# CosGeneGate Selects Multi-functional and Credible Biomarkers for Single-cell Analysis

**DOI:** 10.1101/2024.05.22.595428

**Authors:** Tianyu Liu, Wenxin Long, Zhiyuan Cao, Yuge Wang, Chuan Hua He, Le Zhang, Stephen M. Strittmatter, Hongyu Zhao

## Abstract

Selecting representative genes or marker genes to distinguish cell types is an important task in single-cell sequencing analysis. Although many methods have been proposed to select marker genes, the genes selected may have redundancy and/or do not show cell-type-specific expression patterns to distinguish cell types. Here we present a novel model, named CosGeneGate, to select marker genes for more effective marker selections. CosGeneGate is inspired by combining the advantages of selecting marker genes based on both cell-type classification accuracy and marker gene specific expression patterns. We demonstrate the better performance of the marker genes selected by CosGeneGate for various downstream analyses than the existing methods with both public datasets and newly sequenced datasets. The non-redundant marker genes identified by CosGeneGate for major cell types and tissues in human can be found at the website as follows: https://github.com/VivLon/CosGeneGate/blob/main/marker gene list.xlsx.

## Introduction

Single-cell sequencing technologies offer high-throughput observations into complex biological systems at the cell level ^1,2^, which help elucidate disease mechanisms and improve treatments ^3–5^. These technologies enable the characterization of various molecules, such as DNA (scDNA-seq) ^6^, RNA (scRNA-seq) ^7,8^, and proteins ^9^. They can also facilitate epigenetic studies through single-cell ATAC sequencing (scATAC-seq) ^10,11^ and methylation ^12,13^. There have also been rapid developments of spatial single-cell technologies ^14^. These single-cell technologies have been rated as among the most impactful ones in recent years ^4,15^.

Since single-cell sequencing allows us to study the properties of different cell types, the inference of cell types has become important ^16^. One approach for cell-type annotation is based on marker genes of different cell types ^17,18^. Because scRNA-seq data are high dimensional, sparse and noisy ^19^, there is significant challenge in cell type assignment using marker genes. In addition, genes with similar biological functions tend to show similar expression patterns, further complicating marker gene selections.

Currently, researchers use three sources of information to select marker genes for cell type annotation. The first source is from experts, who select marker genes based on biological knowledge and prior experiments. However, such selections are subjective and lead to different marker gene sets ^20–22^. The second source is through the identifications of differentially expressed genes between groups of cells ^23,24^. By performing the “one-over-all” statistical test for pre-clustered data (using either Louvain ^25^ or Leiden ^26^), these methods select marker genes based on p-values or fold changes or both. However, these methods often assume that the observed expression abundance follows specific distributions, e.g., negative binomial, and test the mean difference between the two groups of cells, which may limit their utilities. Moreover, selecting marker genes based on marginal statistics often leads to genes with similar expression profiles, as shown in Extended Data Figure 1. The third source is through machine learning models ^27,28^, where marker genes are selected based on interpretable machine learning models. A machine learning-based approach uses a dataset with annotated cell types as the training set and trains a classifier with feature selection functions or generates correlation on the training dataset to find the features most relevant to the prediction results of different categories ^29^. Such features are then treated as marker genes. Machine learning-based models can select marker genes as minimal sets with less redundancy. However, the existing machine learning models either fail to capture patterns of different marker genes or consume much computation resource (see the Results section). Moreover, current methods also cannot identify cells with incorrect cell-type annotations, which means the correctness of marker genes may be confused by incorrect cell types of training datasets.

To overcome the limitations of the existing approaches, we present a model based on interpretable neural networks with stochastic **gate** design ^30^ and **cosine similarity** regularization ^31^ for marker **gene** selection, and denote this methods as **CosGeneGate**. Our model selects marker genes with cell-type-specific expression patterns, which are further discussed in the Results section, by utilizing large-scale annotated scRNA-seq data as the training datasets, and it is also scalable for datasets of different sizes with flexible GPU cores. We offer a list of marker genes with their credible sets for major cell types without redundancy of features. Moreover, we consider several downstream applications with the selected marker genes, for example, improving the performance of bulk RNA-seq deconvolution tools, reducing the dimensions of single-cell data, identifying spatially-varying marker genes for spatial transcriptomic data, and refining incorrect cell types. We also illustrate how to apply CosGeneGate to uncover disease-specific marker genes for the immune cells from patients with Alzheimer’s disease.

## Results

### CosGeneGate overview

Given one or more scRNA-seq dataset(s) with rows as samples (cells) and columns as features (genes), and known annotated information (including cell types or cell states), our model aims to learn a projection from a high-dimensional space to a lower-dimensional space that preserves biological information. The subset of genes corresponding to the highest prediction accuracy for a given tissue can be treated as the optimal marker-gene candidates for this tissue. To generate the final list of optimal marker genes, we ensure that these genes can not only achieve the highest cell-type classification accuracy, but also follow the cell-type-specific expression patterns for different cell types supported by previous research ^27,31^.

Our model has two components. The first component is a stochastic gate (STG) neural network, which selects marker genes based on prediction accuracy. The second component is a post-selector based on the cosine similarity of candidate genes generated by STG, which can focus on candidate genes with similar expression patterns. The final gene list is ranked by scores from the second component and filtered to reduce redundancy. When there are no cell labels, we can assign labels from clustering algorithms ^25,26^ or knowledge transformation from another modality ^9^. An overview of CosGeneGate is shown in Figures 1 (a) and (b).

**Fig. 1.**
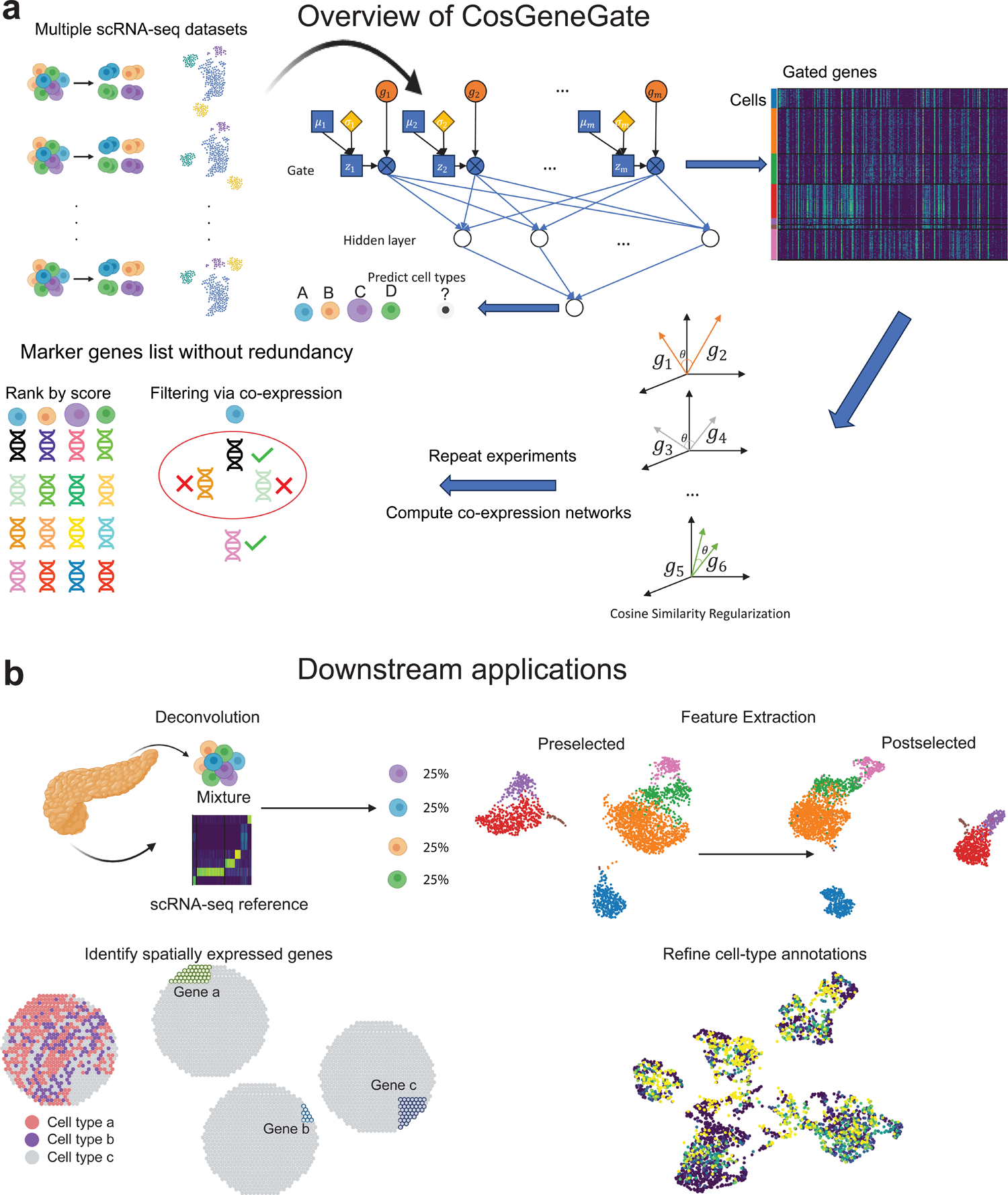
The landscape of CosGeneGate and downstream applications. (a) CosGeneGate utilizes large-scale scRNA-seq as training datasets. Using stochastic gates, we can select candidates of marker genes based on prediction metrics. After collecting all the candidates, we utilize cosine similarity as a method for constraining and filtering target marker genes. Here *g*_1…*m*_represents the input gene expression for *m* genes. For each gene, we have a corresponding gate, denoted as *z*. Each gate is constructed based on a reparameterization trick. The sampling distribution for reparameterization is Gaussian distribution, and *μ* represents the mean and *σ* represents the standard deviation. In the cosine-constrained step, *θ* represents the angle of two gene expression vectors. We run multiple experiments and use correlation to extract equivalent genes, and further filter redundant genes. (b) The downstream applications of marker genes from CosGeneGate include bulk-seq/spatial transcriptomic data deconvolution, scRNA-seq data feature extraction, spatially expressed marker genes identification, and cell-type annotation correction.

We assess the performance of CosGeneGate based on various datasets and metrics in this manuscript. Moreover, we develop a new pipeline for deconvolution analysis to estimate cell-type proportions from bulk RNA-seq datasets.

### Marker genes selected by CosGeneGate improved the performance of cell-type annotation demonstrated by benchmarking analysis

We first investigated the performance of CosGeneGate with three large-scale scRNA-seq datasets from different tissues (PBMC ^29,32–34^, Pancreas ^35^, and Heart ^18^). We illustrate the cell-type similarity across different batches from PBMC in Extended Data Figure 2, as an example to demonstrate the diversity of our training dataset. We compared the prediction accuracy based on marker genes from different models, including genes from experts (expert design), COSG ^31^, NSForest ^27^, scGeneFit ^28^, STG ^30^, and CosGeneGate. By using the leave-one-out strategy, we used each batch as the testing dataset and all other batches of the same tissue as the training dataset for selecting markers and training k-nearest neighbor (kNN) classifiers. The information of batches is provided by the sources of these datasets. We evaluated the performance of all models based on metrics including accuracy, weighted F1 score, label Normalized Mutual Information (label NMI), and label Adjusted Rand Index (label ARI). Figure 2 (a) shows the average scores of the above four metrics for each method and each batch. According to Figure 2 (a), marker genes from CosGeneGate achieved high annotation scores across different datasets. The rank of the annotation scores from CosGeneGate was in the top three for almost all the batches.

**Fig. 2.**
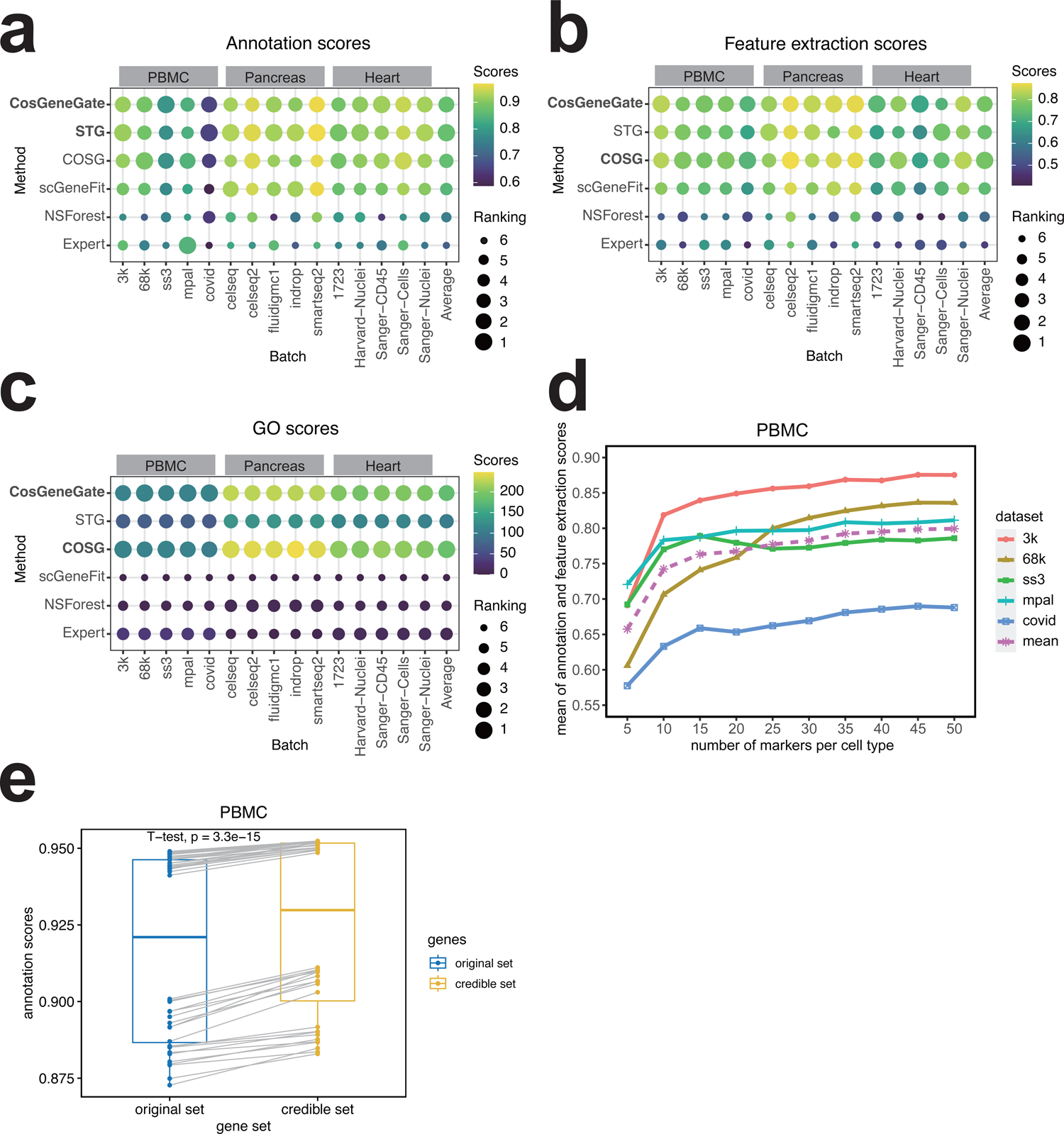
Experimental results for the comparison in the cell-type annotation task. We boldfaced the top 2 methods in panels (a)-(c). (a): bubble plot for annotation scores (i.e. the average score of accuracy, weighted F1, label ARI, and label NMI) for 12 batches of PBMC, Pancreas, and Heart. The last column is the averaged annotation scores for each method. (b) bubble plot for feature extraction scores (i.e. the average of PAGA similarity, scaled ASW, cluster ARI, and cluster NMI) for 12 batches of PBMC, Pancreas, and Heart. The last column is the averaged feature extraction scores for each method. (c) bubble plot for GO scores in 12 batches of PBMC, Pancreas, and Heart. The last column is the averaged GO scores for each method. (d) line chart for hyper-parameter tuning. (e) boxplot and test result for redundancy removal.

Based on the average score of all the datasets, CosGeneGate ranked second among all the competitors, as shown in the last column of Figure 2 (a). Moreover, the variance of annotation scores based on these marker genes is lower than all the other methods except COSG in various metrics, as shown in Extended Data Figures 3-5. COSG does not have a training process, so it is not affected by random numbers. Using marker genes from CosGeneGate is also significantly better than using marker genes from experts for cell-type annotation.

### Validating the marker genes selected by CosGeneGate on the preservation of biological information

Here we investigated the contribution of marker genes for preserving cell-type distinction. By selecting informative marker gene sets (as a method of feature extraction), we can utilize gene expression profiles with a smaller number of features to perform clustering, which is a key step before cell-type annotation. A good marker gene set should preserve the biological distinction induced by cell-type-specific gene expression. Therefore, to evaluate the performance of different gene sets, we computed metrics including PAGA ^36^ similarity, scaled Average Silhouette Width (scaled ASW), cluster ARI, and cluster NMI based on both highly variable genes (HVGs) and marker genes from different methods and averaged them as the final score. Such scores are relevant for the performance of marker genes for clustering and biological preservation. Details of our evaluations can be found in the Methods section. The average scores are summarized in Figure 2 (b). Based on this figure, CosGeneGate outperformed other methods in seven out of the 15 batches we tested. By averaging all batches from all three datasets, CosGeneGate ranked second among all competitors, shown in the last column. Moreover, we computed the Gene Ontology (GO) enrichment results (details are in the Methods section) between each pair of two marker genes and divided the pairs into two groups as genes selected for the same cell type and different cell types. Then, by conducting a t-test between the scores of the two groups, the negative logarithm p-value of the t-test can show if there is a statistically significant difference between genes selected for different cell types, thus showing the abilities of biological information preservation. As shown in Figure 2 (c), CosGeneGate performed better than other methods in the PBMC dataset, while ranked second in Pancreas and Heart. These results demonstrate the ability of CosGeneGate to select functional-specific markers for different cell types. CosGeneGate can also select marker genes with more coherent expressions in the corresponding cell type, and thus these genes have more clear cell-type-specific patterns. Based on Extended Data Figure 6, genes selected by STG are usually expressed across several cell types, whereas COSG usually selects genes with ideal expression profiles, however at the cost of reducing prediction accuracy. Overall, our analysis demonstrated the superiority of CosGeneGate in selecting marker genes for biological information preservation.

Regarding the parameter selection, we used the average score of all annotation and feature selection metrics to select the number of marker genes used in the input of CosGeneGate. Figure 2 (d) shows that 50 is a suitable number of markers for each cell type in the PBMC dataset. Meanwhile, the tuning metrics of the Pancreas dataset are shown in Extended Data Figure 7. For other methods, we tuned their parameters to reach their best performance in each dataset for fair comparison. It is worth noting that, by averaging all evaluation metrics of cell-type annotation and feature extraction from all datasets, CosGeneGate ranked first among all methods, suggesting that marker genes selected by CosGeneGate can optimize these the cell-type annotation and feature extraction tasks simultaneously.

### Removing redundant genes by uncertainty and co-expressions

Here we investigated the performance of CosGeneGate after removing gene redundancy based on changing random seeds and constructing co-expression networks. The workflow of redundancy removal is explained in the Methods section and visualized in Figure 1 (a). We compared the four metrics of cell-type annotation between the original marker gene set and the credible marker gene set (i.e., the gene set after redundancy removal). For the PBMC dataset, the result of the paired t-test between the two gene sets is shown in Figure 2 (e), and the other results are shown in Extended Data Figure 8. According to the boxplot and test result, we improved annotation accuracy for PBMC and Pancreas by generating a credible set of genes, demonstrating that removing redundant genes can lead to more accurate cell-type annotation. We have provided credible sets for each major cell type of different tissues (PBMC, Pancreas, and Heart) in Supplementary file 1.

### Explainable gene expression patterns of CosGeneGate

In this section, we investigate the preference of CosGeneGate for marker gene selection and present an algorithm to choose a suitable number of marker genes from a statistical perspective. We computed the Wilcoxon rank-sum test score for the marker genes from PBMC and Pancreas based on candidate genes selected by CosGeneGate, and visualized the relation between the number of marker genes and the score in Extended Data Figure 9 (a). This figure shows that the number of marker genes with the best performances depends on the most difficult cells to classify in the tissue, i.e., the cells with the highest number of genes needed for the score to reach its maximum value. For example, in the PBMC dataset, selecting 50 genes per cell type is the optimal choice based on our metrics. On the other hand, in the Pancreas dataset, selecting 20 genes per cell type is the optimal choice based on our metrics. These two choices correspond to the cell types highlighted by the red block in Extended Data Figure 9 (a). We also tried to use two different approaches to explain the preference of our marker genes, known as the COSG score (shown in Extended Data Figure 9 (b)) and the Metamarker score ^37^ (shown in Extended Data Figure 9 (c)). However, in the analysis of the COSG score for the PBMC dataset, we did not see a positive relation between the COSG score and the performance of marker genes. The original manuscript for Metamarker score advocates to choose 50-200 genes per cell type for accurate cell-type annotation based on the trade-off between the fold change (FC) score and the area-under-curve (AUC) score. However, based on our analysis, not all the cell types had such patterns, and it is difficult to balance between the FC score and the AUC score for all cell types in the Pancreas dataset. Therefore, previous methods could not be used to explain the multifunctional marker genes from CosGeneGate, and our analysis is more suitable for model explanation based on the statistical test. Details of our algorithm can be found in the Methods section.

### Selecting marker genes improved the performance of deconvolution

We first investigated the impact of marker gene selection on deconvolution. For signature matrix-based deconvolution methods, we can deconvolve the bulk RNA-seq datasets by only keeping selected marker genes rather than using all genes. Since only informative genes are kept, the deconvolution performance is expected to be better. Then, we compared the deconvolution results of all genes, NSForest ^27^, scGeneFit ^28^, STG ^30^ and COSG ^31^ on different deconvolution models: CIBERSORTx ^38^, MuSiC ^39^, and NNLS ^40^. To evaluate the performance, we used Root Mean Square Error (RMSE), Pearson Correlation Coefficient (PCC), and Coefficient of Variation (CV) as metrics. Details of these metrics are included in the Methods section.

Regarding the parameter selection, we used RMSE to select the optimal number of marker genes. For other competitors, we tuned the number of marker genes to reach their best performance accordingly for a fair comparison. Empirical experiments show that the optimal number of marker genes differs for different datasets and different gene selection strategies. Therefore, it should be tuned in practice. To tackle the problem, we designed a deconvolution pipeline that adjusts the number of marker genes to be selected by CosGeneGate. Figures 3 (a) and (b) show the overall workflow of the pipeline for the pseudo bulk mode and the real bulk mode, respectively. In the pseudo bulk mode, the scRNA-seq dataset is split into two parts, one for pseudo bulk data ^38^ generation, and the other for hyperparameter tuning with CosGeneGate marker genes on these generated pseudo bulk mixtures for this model. In the real bulk mode, the model is directly tuned on the real bulk data ^41^, and its ground truth cell-type proportion information is provided by the users. Figure 3 (c) shows how the deconvolution performance varies as the number of marker genes for each cell type changes within the pancreas dataset. There is a parallel pattern for RMSE between the validating dataset and the testing dataset, suggesting that the number of marker genes that achieve the best performance in the validation dataset should also achieve almost the best performance in the testing data, guiding the selection of the number of marker genes. Figure 3 (d) shows the performance of deconvolution on a real bulk dataset known as the 3celllines dataset. CosGeneGate performed well on average compared with other gene selection strategies, and outperformed the case of choosing all genes. Figure 3 (e) shows the performance improvement for both the pseudo bulk mode and the real bulk mode. We can observe that in most cases, the pseudo bulk mode improves the performance of deconvolution, which is further improved by the real bulk mode.

**Fig. 3.**
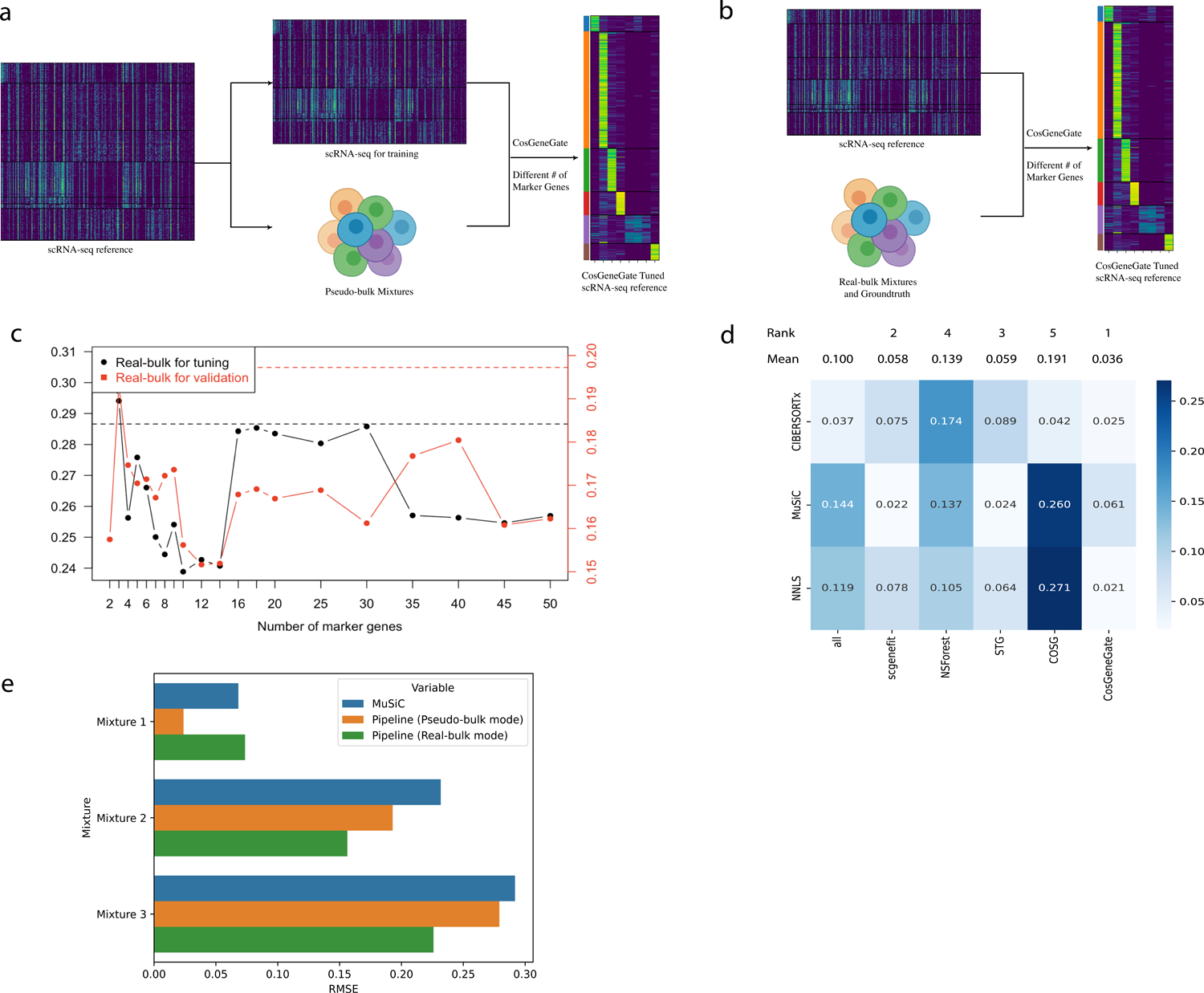
Experimental results for the comparison in the deconvolution task. (a) The workflow for CosGeneGate marker-based deconvolution pipeline (Pseudo-bulk Mode) (b) The workflow for CosGeneGate marker-based deconvolution pipeline (Real-bulk Mode) (c) Relationship between the number of markers and deconvolution performance (d) Benchmark of performance of deconvolution in the real bulk three-cell dataset (e) The performance of deconvolution pipeline in real bulk pancreas dataset with two different modes.

### Selecting marker genes uncovered spatially-informed patterns in spatial transcriptomic data

We next assessed the performance of CosGeneGate on selecting marker genes from scRNA-seq data for spatial transcriptomics studies. We considered two types of spatial transcriptomic data to investigate the general applicability of our method. In the dataset (visium_fluo) sequenced using 10x Visium ^42^, every spot within the spatial transcriptomic dataset encompasses a variable number of cells and 10x Visium is a whole-transcriptome-based technique. In contrast, in the dataset (human breast) obtained through Xenium sequencing ^43^, we can extract a single-cell gene expression profile with hundreds of genes, and each spot corresponds to an individual cell. Using scRNA-seq to select marker genes for the spatial transcriptomic data is meaningful for spatial-level deconvolution and cell-type annotation ^44^. We compared the results of 2000 highly variable genes (HVGs), COSG, NSForest, scGeneFit, scMAGS ^44^, STG, and CosGeneGate to evaluate their abilities to identify marker genes for spatial transcriptomic data from scRNA-seq data. We added scMAGS in this section because of its ability to analyze spatial transcriptomic data. Figure 4 (a) shows the evaluation of the preservation of gene-gene correlation selected by different methods for the 10x Visium dataset and the human breast dataset based on different random seeds. According to Figure 4 (a), CosGeneGate has a high gene-gene correlation across different methods and different datasets. Using marker genes from CosGeneGate is also significantly better than using HVGs. Details of our evaluation methods are summarized in the Methods section. Moreover, Figures 4 (b) and (c) show the distribution of cortex of the visium_fluo dataset and cell types of the human breast dataset with spatial location respectively. The gene expression profiles of spatially-expressed marker genes shown in Figure 4 (d) of malignant cells identified by CosGeneGate can illustrate a binary expression profile compared with cells from different cell types of the human breast dataset. These results altogether support the ability of CosGeneGate to select marker genes that uncover cell-type-specific gene expression patterns in spatial transcriptomic data.

**Fig. 4.**
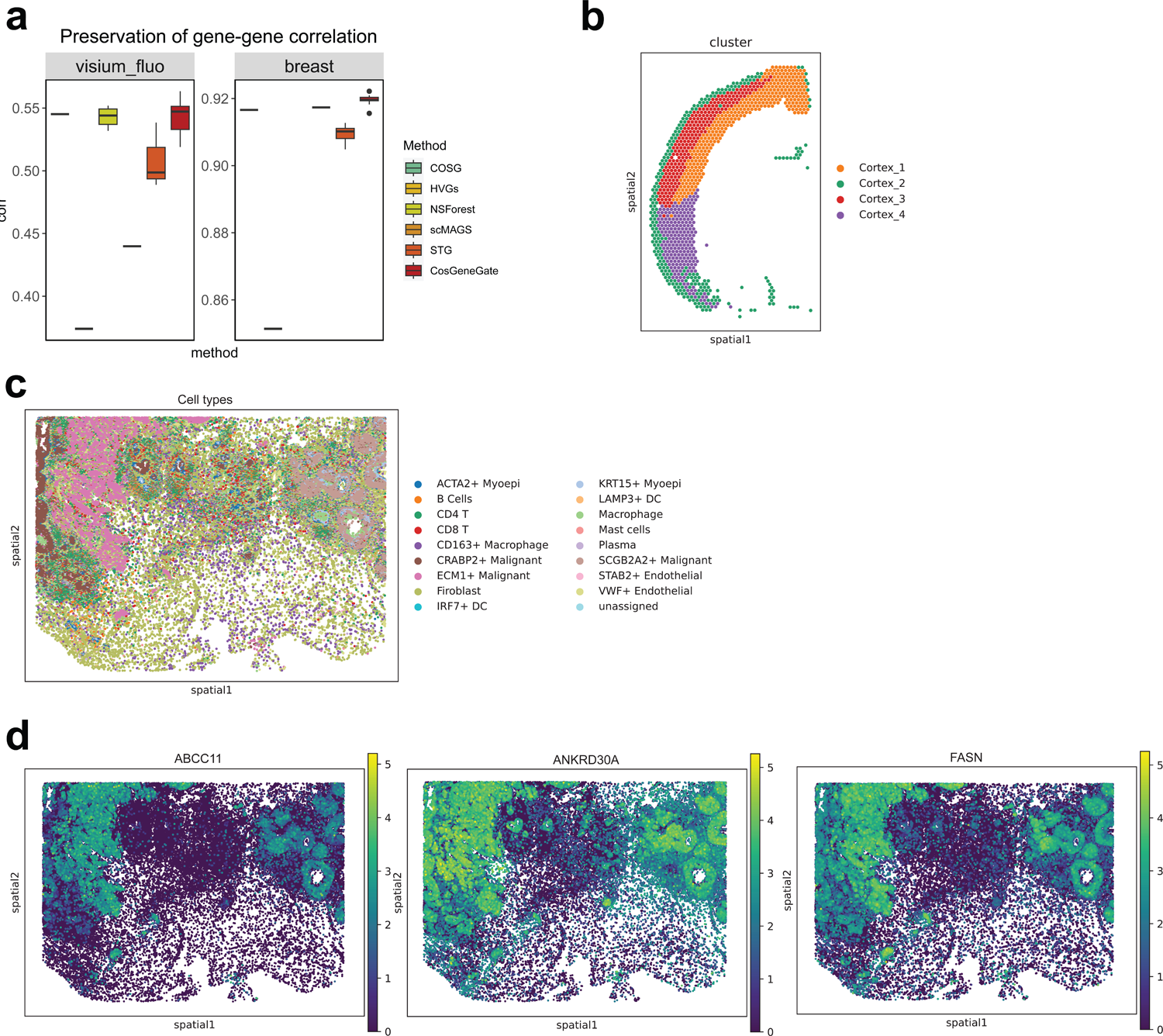
Experimental results for the application of marker genes. (a) The evaluation for the preservation of gene-gene correlation of the genes selected by different methods for the 10x Visium dataset and the human breast dataset. We did not report the results of NS-Forest for the human breast dataset because its running time exceeded our limit. (b) Visualization of cluster distribution of the 10x Visium dataset. (c) Visualization of the cell-type distribution of the human breast dataset. (d) Spatially-expressed marker genes identified by CosGeneGate in the human breast dataset.

### Selecting marker genes from CosGeneGate refined uncertain cell types

We assessed CosGeneGate’s ability to use selected marker genes to detect incorrect or unknown cell types. According to Figure 5 (a), the 3-fold intra-dataset hierarchical classification accuracy of all methods is low, leading us to conjecture that there are wrongly labeled sub-cell types in the Zeisel dataset ^45^. Figure 5 (b) shows the UMAP for the sub-cell type distribution of the Zeisel dataset. In Figure 5 (c), we count and then display the number of times a cell’s classification result did not match the label divided by the total number of trials as the uncertainty score. As can be seen in the two UMAPs, tightly clustered cells of the same cell type tend to have uncertainty scores closer to zero and cells between cell-type clusters have higher uncertainty scores, which accords with intuition. We defined unknown or known sub-cell types as wrongly or correctly labeled sub-cell types by markers from CosGeneGate using all ten seeds and all ten n_neighbors parameters of kNN classifiers, i.e., cells’ uncertainty scores equal zero for cells which are classified as their annotated types under all the experiments. Then, using the expert markers from Zeisel ^45^, we generated the dotplots of unknown and known sub-cell types in oligodendrocytes, as shown in Figure 5 (d). Comparing the upper panel with the bottom panel, we can observe that the expert marker genes from unknown cell types such as Oligo 5 and Oligo 6 have unclear patterns. It can be seen that the cells with unknown labels are indeed wrongly labeled based on their expression of the expert markers. We offer the full marker gene list in Extended Data Figure 10. Finally, we collected the experiment results from the kNN classifier and re-annotated cell types with the maximal cell-type proportion of prediction. These results are summarized in Figure 5 (e).

**Fig. 5.**
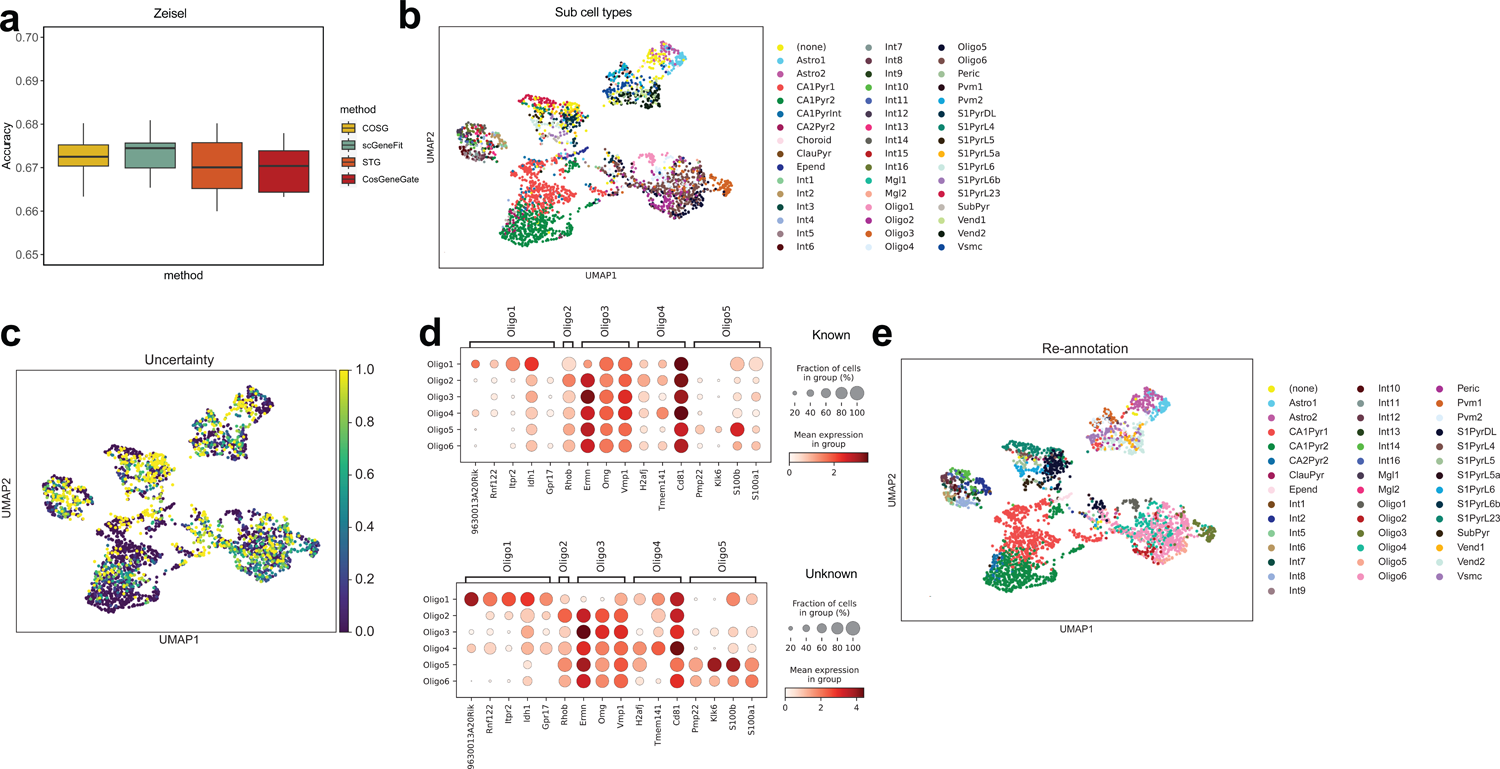
Utilizing marker genes to detect unknown cell types. (a) The accuracy of cell-type annotation based on different methods for the Zeisel dataset. (b) UMAPs for the sub-cell type distribution of the Zeisel dataset. (c) Sub-cell type-specific gene expression plots for cells with incorrect labels (top) and cells with correct labels (bottom). (d) UMAPs for the cells with uncertainty. (e) UMAPs for the re-annotated cells.

### Discovery of disease-specific marker genes for Alzheimer’s disease

CosGeneGate can also discover disease-specific marker genes based on the analysis of sub-cell types across samples with different conditions. Here we focus on Alzheimer’s disease (AD), the most common age-related neurodegenerative disease. Clinical symptoms of AD are characterized by progressive cognitive decline and dementia. Therefore, the analysis of possible genetic risk factors for AD is important for clinical practice today. Microglia cells are important immune cells in the brain, which are highly correlated with AD ^46–49^. To identify disease-specific marker genes, we sequenced a new dataset containing 126,687 cells from seven samples using 10x Multiome sequencing. We show the distribution of diseased conditions in Figure 6 (a) and the distribution of cell types in Figure 6 (b). Here we tried three different approaches and used the overlap score (known as precision) between selected disease-specific marker genes and known disease-associated genes as the criterion. Details are included in Extended Data Figure 11. Finally, we identified the sub-clusters/sub-cell types of microglia cells from our AD-Health Control (HC) datasets and then utilized CosGeneGate to select possible genetic risk factors. Such genes were identified in the AD samples as marker genes of certain sub-cell types rather than in HC samples. Figure 6 (c) shows the expression patterns of selected disease-specific marker genes. The selected AD-specific marker genes showed higher expression levels in AD-associated sub-cell types. The overlapping information between selected marker genes and known AD-associated genes can be found in Supplementary file 2. Those genes which are not previously identified as AD-associated can be treated as novel AD-specific marker genes. Therefore, we can also use CosGeneGate to identify the risk genes for diseases supported by disease-specific expression patterns and the results for mining AD-associated genes from known databases.

**Fig. 6.**
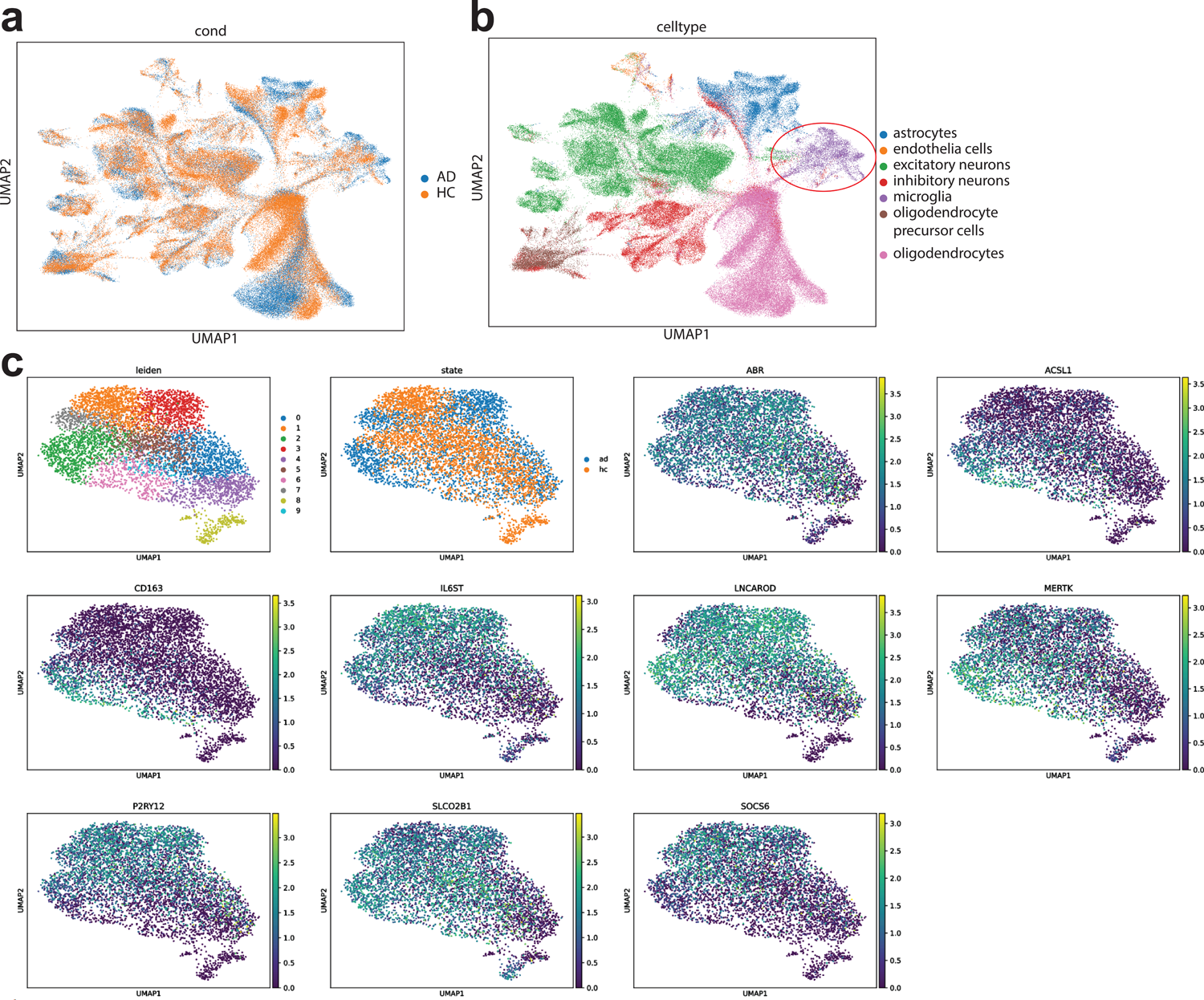
Utilizing marker genes for sub-cell types to explore the biological system in the human brain. (a) UMAPs of the AD-HC dataset colored by disease condition. (b) UMAPs of the AD-HC dataset colored by cell type. Microglia cells are highlighted by a red circle. (c) UMAPs for the AD-specific marker genes of Microglia cells.

## Discussion

Marker genes not only work as the safeguards for the cell-type annotation prior to downstream analysis of single-cell data, but can also facilitate downstream applications including bulk RNA-seq data deconvolution, feature extraction, spatial gene expression pattern identification, sub-cell type correction, the discovery of disease-specific genes, and others. Therefore, the multifunctional marker genes selected by CosGeneGate are important for single-cell data pre-processing and downstream applications. CosGeneGate selects marker genes by considering both the contribution of target genes to cell-type annotation, and the cell-type-specific expression patterns of marker genes. By combining these two ideas together, we showed the superiority of marker genes identified by CosGeneGate in the Results section. Furthermore, we designed a framework to reduce the redundancy of our marker gene list and demonstrated its usefulness in the Results section.

We note that in the cell-type annotation task, CosGeneGate had similar performance compared to STG and COSG. However, by taking a deeper look at the gene expression patterns of STG, the marker genes selected by STG tended to express across different cell types, which were false-positive signals discovered by STG shown in Extended Data Figure 6. Moreover, the genes selected by COSG are not ideal choices for other downstream applications, including representative feature extraction and bulk RNA-seq deconvolution. It also consumed more memory usage compared to CosGeneGate. Our method integrated the advantages of COSG and STG while avoiding their shortcomings. CosGeneGate also outperformed NSForest and scGeneFit for multiple tasks. The running time of NSForest was too long to select marker genes effectively. Genes selected by scGeneFit are not cell-type-specific and do not have rank information to quantitatively describe their quality, which limits their functionality.

For the deconvolution task, we developed a pipeline for bulk RNA-seq data deconvolution, which allowed users to choose different modes and it could select the number of marker genes with the best performance automatically, which is also a user-friendly design. For the feature extraction task, we demonstrated that marker genes selected by CosGeneGate could not only preserve the distinction of cell types, but also retain the structure similarity of different cell types before running the selection step. Reducing the dimensionality we need for single-cell analysis is important for efficiency. For the analysis related to spatial transcriptomic data, we demonstrated that marker genes selected by CosGeneGate from scRNA-seq datasets could be used to identify genes with spatial expression patterns based on the spatial transcriptomic data from the same tissue, which builds a bridge for multi-omics data analysis. For the function related to cell-type correction, we used CosGeneGate to refine the sub-cell types from scRNA-seq datasets thus increasing the data quality. Therefore, the marker genes from CosGeneGate are high-quality across different scenarios.

We also used CosGeneGate for biological discoveries, for example, identifying new disease-specific marker genes based on sequenced datasets. We identified a number of disease-specific marker genes for the sub-cell types of Microglia cells by analyzing samples with/without AD. In the final marker gene set, some genes are verified as disease-associated markers by analyzing GWAS statistics and/or related biological experiment results. The rest of genes can be treated as newly discovered marker genes. Therefore, the application of CosGeneGate towards disease analysis is also promising.

To determine the number of marker genes identified by CosGeneGate, we ran experiments by adjusting the number of marker genes as a hyper-parameter and selecting the number corresponding to the cell-type annotation and feature extraction task. We also considered the explainability of our methods and discovered the relation between the number of marker genes per cell type and the related score from statistical tests. After redundancy removal and co-expression patterns, we summarized these genes into a table for major tissues, which could be used by users for their specific tasks. Moreover, the number of marker genes is also an editable hyper-parameter. Users can also adjust its value if they prefer more/fewer markers.

In summary, CosGeneGate can select high-quality marker genes for multiple downstream applications and has acceptable running time and memory usage. It has its unique feature selection design by combining ideas from the machine learning area and the biology area. The next step of CosGeneGate will shift to multi-omic data analysis and multi-species data analysis. We also plan to use CosGeneGate to discover the relation among genes, peaks, proteins, as well as similar genes across different species.

## Methods

### Problem definition

In the marker gene selection problem, we denote the scRNA-seq expression profile as a matrix *X*^*N*×*p*^, where *N* represents the number of cells, and *p* represents the number of genes. We use *Y*^*N*×1^ to represent ground truth cell types. Based on CosGeneGate, our target is to find an optimal set of genes, which can minimize the loss in our training process and maximize the mutual information between cell types and marker genes. Therefore, we have:

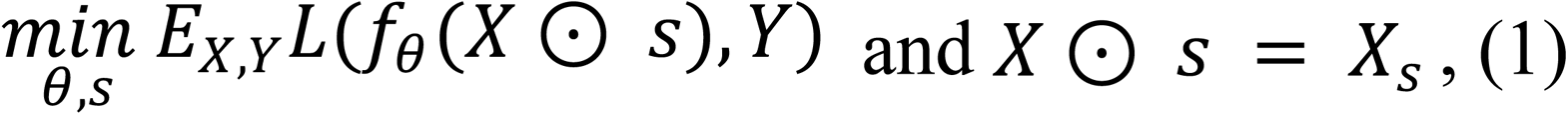

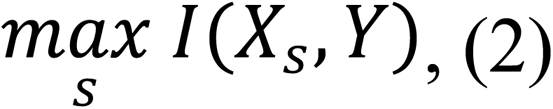

where *L*(⋅) represents the loss function, and *f*_*θ*_(⋅) represents the CosGeneGate model. Here *s* denotes the indices of selected marker genes, and *s* ⊂ {1,2, …, *p*}. *X*_*s*_ represents the optimal marker gene sets. We use *I*(*X*_*s*_; *Y*) = *H*(*Y*) − *H*(*Y*|*X*_*s*_) to represent the mutual information between the set of marker genes and the set of cell types. *H*(*X*_*s*_) denotes the entropy of distribution *p*_Y_ and *H*(*Y*|*X*_*s*_) denotes the entropy of distribution *p*_Y|*Xs*_. We also control the number of selected marker genes for different cell types, |*s*| = *Ka*, where *K* represents the number of cell types and *a* represents the number of marker genes for each cell type.

### CosGeneGate

CosGeneGate is designed to find a marker gene set with high prediction accuracy and also preserving biological information. The main idea of CosGeneGate is to first select a subset of genes with higher prediction accuracy, and then utilize cosine similarity to select the final marker gene set from the subset. It consists of several steps, which are detailed in the following.

The first step is a stochastic gate (STG) neural network, which utilizes a *l*_0_ norm and Bernoulli gates applied to each of the input nodes of a neural network. The number of gates is *D*. We denote each Bernoulli gate as a random vector *S*_*d*_, whose entries are independent and identically distributed, satisfying *P*(*S*_*d*_ = 1) = *π*_*d*_ for *d* ∈ {1,2…, *D*}, respectively. Then problem (1) becomes:

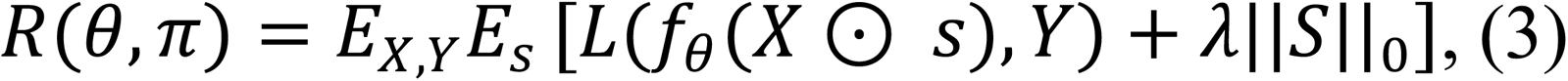

where we denote the empirical expectation over the observations as 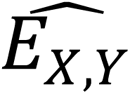, and due to the Bernoulli distribution of S, we have 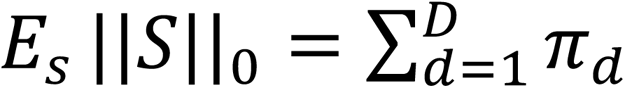. Thus, the problem of feature selection is simplified as finding the optimal Bernoulli distribution parameters, *θ* and *μ*, that minimize the empirical risk shown in Eq. 4 in the following. Then we utilize a Gaussian-based continuous relaxation that is fully differentiable and only activates or deactivates gates linking each feature (node) to the rest of the network. With *γ*_*d*_drawn from *N*(0, *σ*^2^) and *σ* fixed through training, each relaxed Bernoulli variable is denoted as sa stochastic gate (STG), which is defined by *z*_*d*_ = *max*(0, *min*(1, *μ*_*d*_ + *γ*_*d*_)). The feature selection problem becomes:

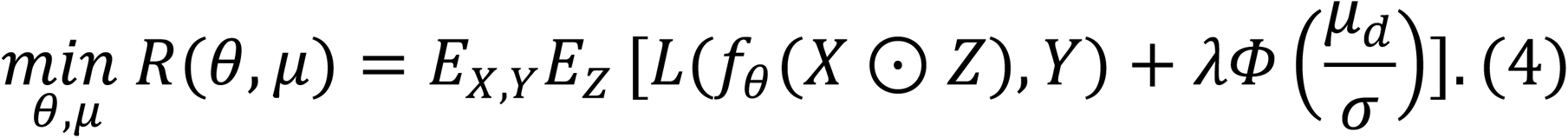

Differentiating with respect to *μ* and utilizing a Monte Carlo sampling gradient estimator, we have 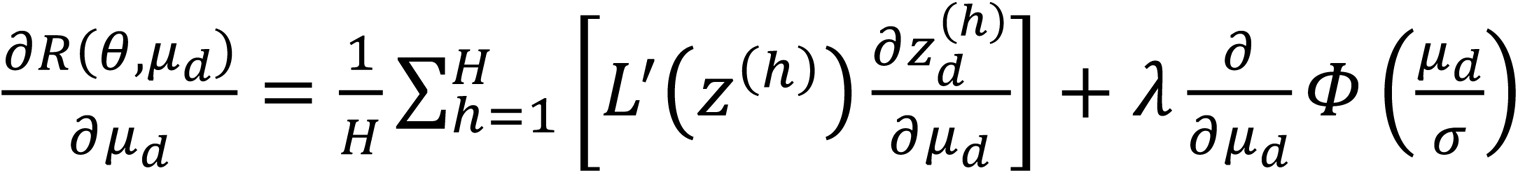, where H is the number of Monte Carlo samples. Therefore, we can update *μ*_*d*_ using the gradient descent for each *d* ∈ {1,2… *D*}. Optimizing Eq. 4, we can get the probability of whether each feature should be selected. Then, ranking each gene with the probability, we can get a list of candidate genes, default as 200 genes. Users can also define a threshold for the probability and select all genes above the threshold.

The second step is a post-selector based on the cosine similarity of candidate genes generated by STG. After the first step, we can get the expression matrix of candidate genes, denoted as *X*^*N*×*M*^, where N is the number of cells and M is the number of genes selected by STG. Refer to K as the number of cell types and *G*_*k*_, *k* ∈ {1,2… *K*}, as cells of cell type k predefined by manual annotation by biological experts or unsupervised clustering, we generate an ideal marker gene *λ*_*k*_ for *G*_*k*_: *λ*_*k*_ = [*λ*_1*k*_, *λ*_2*k*_, … *λ*_*nk*_]′, where *λ*_j*k*_ = 1 if the *j*^*th*^ cell *c*_j_∈ *G*_*k*_ and *λ*_j*k*_ = 0 if *c*_j_∉ *G*_*k*_. Then, we can calculate the cosine similarity between the i^th^ gene *g*_*i*_ and the ideal marker gene *λ*_*k*_ for cell type k as:

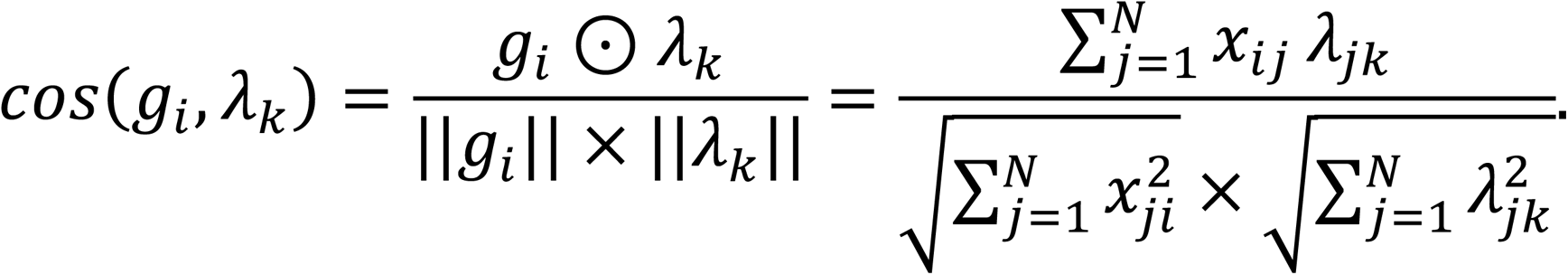

Therefore, based on the cosine similarity and given a penalty factor *τ* (*τ* ≥ 0 and default as 1) for expression of target gene in non-target cell type, the score for evaluating each gene is:

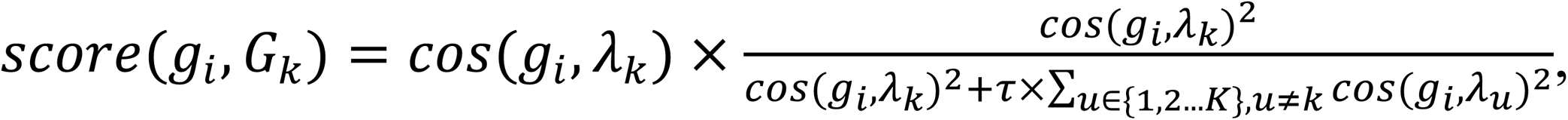

where a higher score represents a higher probability of *g*_*i*_ as a marker gene of cell type k. Therefore, we can rank the candidate genes selected by STG and select the top *a* (*default as* 50) genes as the final marker list.

### Select marker genes without redundancy

According to Figure 2 (e) and Extended Data Figure 7, the uncertainty or randomness does affect the performance of CosGeneGate in performing cell-type annotation or discerning different cell types. Therefore, we consider using this property to remove redundant genes in our post-selection marker gene sets. We also use the genes after removing redundancy to formalize a marker gene list for different cell types across different tissues. By changing the random seeds and running 1,000 experiments for one dataset, we collect the number of times each marker gene was selected for each cell type in the experiments. Here we choose the marker genes that are included in all our experiments as the optimal marker gene sets. This restriction means that we should not remove these genes in our downstream analysis, referred to as optimal marker gene sets ^50^. If there are no markers that are selected in all 1,000 experiments, we use the genes that are included the greatest number of times as the optimal marker gene sets.

For each cell type, we use CS-CORE ^51^ to infer the co-expression relation between the optimal marker genes and the other selected marker genes. The initial credible gene set of each cell type are the optimal marker gene sets. Then we implement the credible set selection method as an iterative two-step process of within and cross cell types: (1) Within each cell type, we select the gene pair with the lowest average co-expression score with each gene in the credible set; (2) comparing among different optimal genes and different cell types, the gene pair with the lowest average co-expression score is selected into the credible set.

This iteration stops until the total number of markers selected for each cell type equals the optimal number of marker genes after parameter tuning. Therefore, we can generate a list containing marker genes without redundancy for each cell type. By repeating this process for different datasets, we offer the cell-type-specific and non-redundant marker gene list for different tissues.

### Interpretation of marker genes selected by CosGeneGate

To explain the patterns of genes selected by CosGeneGate and link the number of markers to the performance of CosGeneGate on cell-type annotation and feature extraction tasks, we propose a method based on candidate selection and statistical tests.

Considering a given dataset *X*^*N*×*p*^, we have *K* cell types per dataset and *a* marker genes per cell type. We first filter the genes of the given dataset and ensure the set of genes only contain selected markers for all cell types. Therefore, we have *X*′^*N*×*Ka*^(by assuming no overlap marker genes for different cell types). We then run the Wilcoxon rank-sum test using *X*′ for each cell type, and we have a list of normalized z-score *W* for every gene in every cell type. We compute the average score 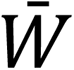 of marker genes for different cell types, and then we compute the 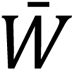 for different numbers of marker genes.

Therefore, we can use 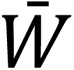 to explain the marker genes selected by CosGeneGate statistically. Moreover, we can analyze the relationship between the number of marker genes and the performance of CosGeneGate by analyzing the relationship between the number of marker genes and the average score.

### Deconvolution pipeline

The deconvolution pipeline for CosGeneGate is designed to identify the signature matrix with the best performance to be used in deconvolution. The main idea is to test deconvolution for different numbers of marker genes |*s*| in a given set *X*_*s*_, then choose the number of marker genes which has the best performance over a given metric. The number of marker genes with the best performance based on RMSE |*s*|_*o*_ can be then calculated as:

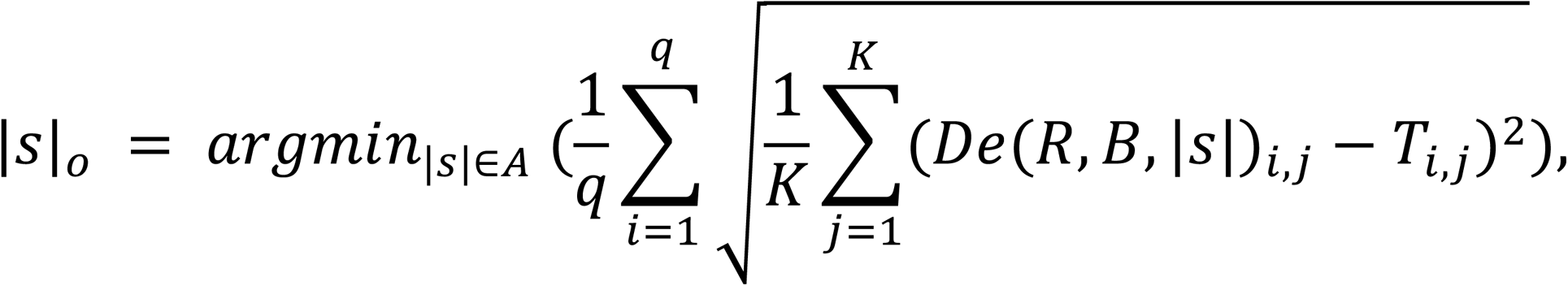

where K is the number of cell types in the mixture, *q* is number of mixtures to deconvolve, *De* is the deconvolution model, R is the single cell reference matrix containing all genes, *B* is the bulk mixture to be tested on, *T* is a ground truth matrix for the bulk mixture *B*, and *A* is the set of potential number of marker genes.

Next, we are able to obtain the marker gene set with the best performance *M*_*o*_ used in deconvolution. Let *M*(*G*_*k*_) denote the marker gene lists for CosGeneGate of cell type *G*_*k*_, *k* ∈ {1,2… *K*} in a descending order of *score*. Then *M*_*o*_ can be written as the union of the top |*s*|_*o*_ marker genes for each cell type *G*_*k*_, i.e.

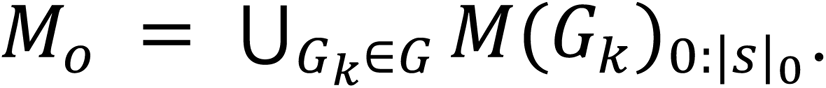

As a result, the pipeline gives a fine-tuned single cell signature matrix *R*_*o*_by only containing the selected marker genes in set *M*_*o*_. In other words, 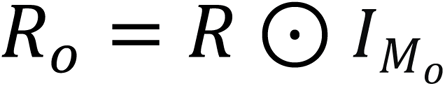 where 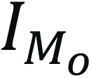 is the identity matrix where only marker genes *g* ∈ *M*_*o*_ is 1.

Depending on the specific dataset users have, the CosGeneGate deconvolution pipeline consists of two modes: pseudo bulk mode and real bulk mode.

For the pseudo bulk mode, users only have to give a single cell reference matrix *S* as input. Then the reference matrix *R* will be split into two parts, *R*′ as the reduced signature matrix and *RP* as the single cell profile to generate pseudo bulk mixtures. The default proportion of this split is *prop* = 0.9, which means 90% for *R*′ and 10% for *RP*. Note that the proportion needs to be selected with caution here, because a small *prop* will significantly reduce the size of the signature matrix, and a signature matrix with less information is typically more difficult to improve the performance although the number of marker genes is fine tuned. On the other hand, if *prop* is too close to 1, there will not be enough information for pseudo bulk to be generated meaningfully and thus make the fine tune process less efficient. Once *RP* is obtained, we can generate the signature matrix by randomly sampling each cell in *RP*, then aggregating the selected cells together to generate the pseudo bulk mixtures *B*, i.e., 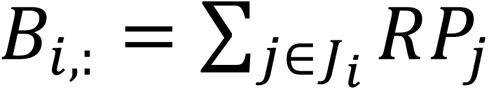,: where *J*_*i*_ is the randomly selected cells in the *i*^*th*^ random mixture. When aggregating cells to be pseudo bulk mixtures, the number of different cell types is also recorded to calculate the proportion of different cell types in each mixture. In this way, the ground truth *T* is obtained, and we are able to find out the best number of marker genes *n*_*o*_ for this dataset.

As for the real bulk mode, users must give a single cell reference matrix *R*, a real bulk mixture *B* and the ground truth *T* for the real bulk mixture *B* as input. Then the best number of marker genes |*s*|_*o*_ can be calculated directly using the given information. The real bulk mode typically requires more information from the user. Users may give the real bulk data and the corresponding ground truth themselves. The sources of ground truth information can come from a different sample in the same tissue but with known cell-type proportion. They can either search for the real bulk data and its ground truth on some existing datasets which may be similar to the mixtures they want to deconvolve. Real bulk mode is recommended over the pseudo bulk mode because there is no need to sacrifice the original signature matrix *R*. It also gives a more reasonable best number of marker genes |*s*|_*o*_ as real bulk is better in practice than artificially generated pseudo bulks.

### Disease-specific marker gene selection

Our first step is to identify marker genes for microglia cells for AD and healthy samples. The second step is to find marker genes that are only identified in the AD samples by removing the common marker genes. After obtaining such genes, we re-cluster the microglia cells from AD samples and run CosGeneGate to identify disease-specific marker genes for sub-cell types. This method avoids possible noise brought by the batch effect. We visualized the expression levels of these genes based on combining AD samples and healthy control samples.

We investigated the functions of the selected genes based on both GWAS summary statistics ^52^ for AD-associated genes and biological experiments. For genes without such information, we treated them as novel disease-specific marker genes. All the genes identified by CosGeneGate are summarized in Supplementary file 2.

## Metrics

### Marker genes selection

To evaluate the quality of selected marker genes, we use a k-nearest neighbor (kNN) classifier, with the parameter of n_neighbors ranging from 5 to 14, to predict the cell types in the testing dataset with marker genes selected from the training datasets. We chose to use the kNN classifier because it does not require further training and it has a better performance compared with Support Vector Machine (SVM) with rejection, which is the best method for cell-type classification supported by ^53^, shown in Extended Data Figure 12. We denote the predicted cell types as 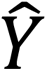 and ground truth cell types as *Y*, the metrics accuracy and weighted F1 scores are defined below:

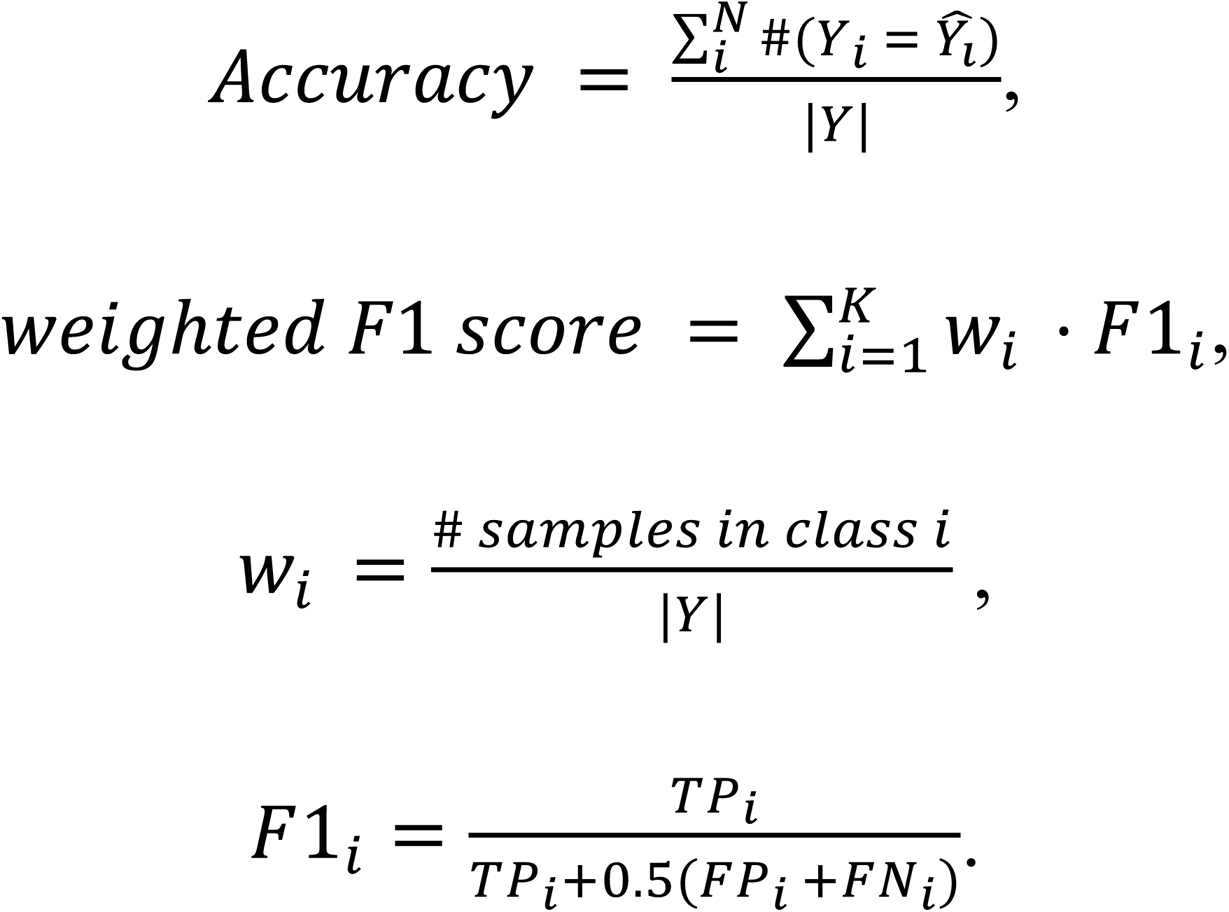

Here *TP*, *FP*, *and FN* denote the true positive rate, false positive rate, and false negative rate, respectively. The final score is the average score of kNN classifiers. Higher accuracy or weighted F1 score represent better marker genes. We also calculate label ARI and label NMI for the cell-type annotation task based on scikit-learn ^54^.

To evaluate the performance of marker genes in feature extraction, we also consider different metrics. To consider the level of cell-cell similarity preservation, we use PAGA ^36^ similarity as a metric. We first compute the topological similarity between the expression profile before marker gene selection and after marker gene selection. We then calculate the difference between these two matrices and smooth the result using a Gaussian kernel. Moreover, we also use Normalized Mutual-information (NMI), Adjusted Rand Index (ARI), and Average Silhouette Width (ASW) of cell types as three metrics to evaluate the quality of marker genes based on scIB ^55^. Higher scores represent better biological information preservation based on selected marker genes.

Since the selected marker genes should be representative of one cell type, we also consider evaluating the quality of marker genes based on gene-gene interaction in one cell type or across different cell types. Here we develop a new metric based on Gene Ontology Analysis (GO Analysis) ^56^ to evaluate the functions of selected genes for preserving cell-type specific information. Such a metric is known as a test for cell-type specific functional genes. Considering we have *m* cell types, we select *k* (*k* ≥ 1) genes for each cell type. We compute the gene-gene similarity based on the Jaccard similarity of the GO enrichment pathways information ^57^ for two genes. The Jaccard similarity for two sets *⍺* and *β* is defined as:

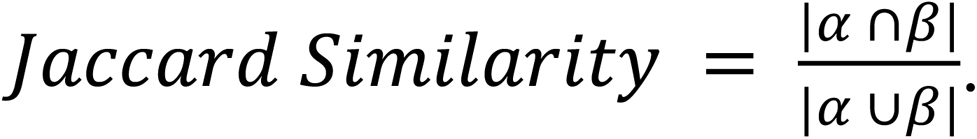

Therefore, we have two groups of Jaccard similarity scores, one denotes the score list for marker genes in the same cell type, while the other one denotes the score list for marker genes across different cell types. Therefore, our null hypothesis *H*_0_is that selected genes do not have cell-type specific functions. Our alternative hypothesis *H*_1_is that selected genes have cell-type specific functions. We run a t-test for the verification step and record the p-value after the negative logarithm transformation. A higher transformed p-value represents a better marker gene set.

### Deconvolution

To evaluate the deconvolution for cell types, we consider the root mean square error (RMSE), mean absolute error (MAE), Pearson’s correlation coefficient (PCC), and coefficient of variation (CV). All of the metrics can be summarized as:

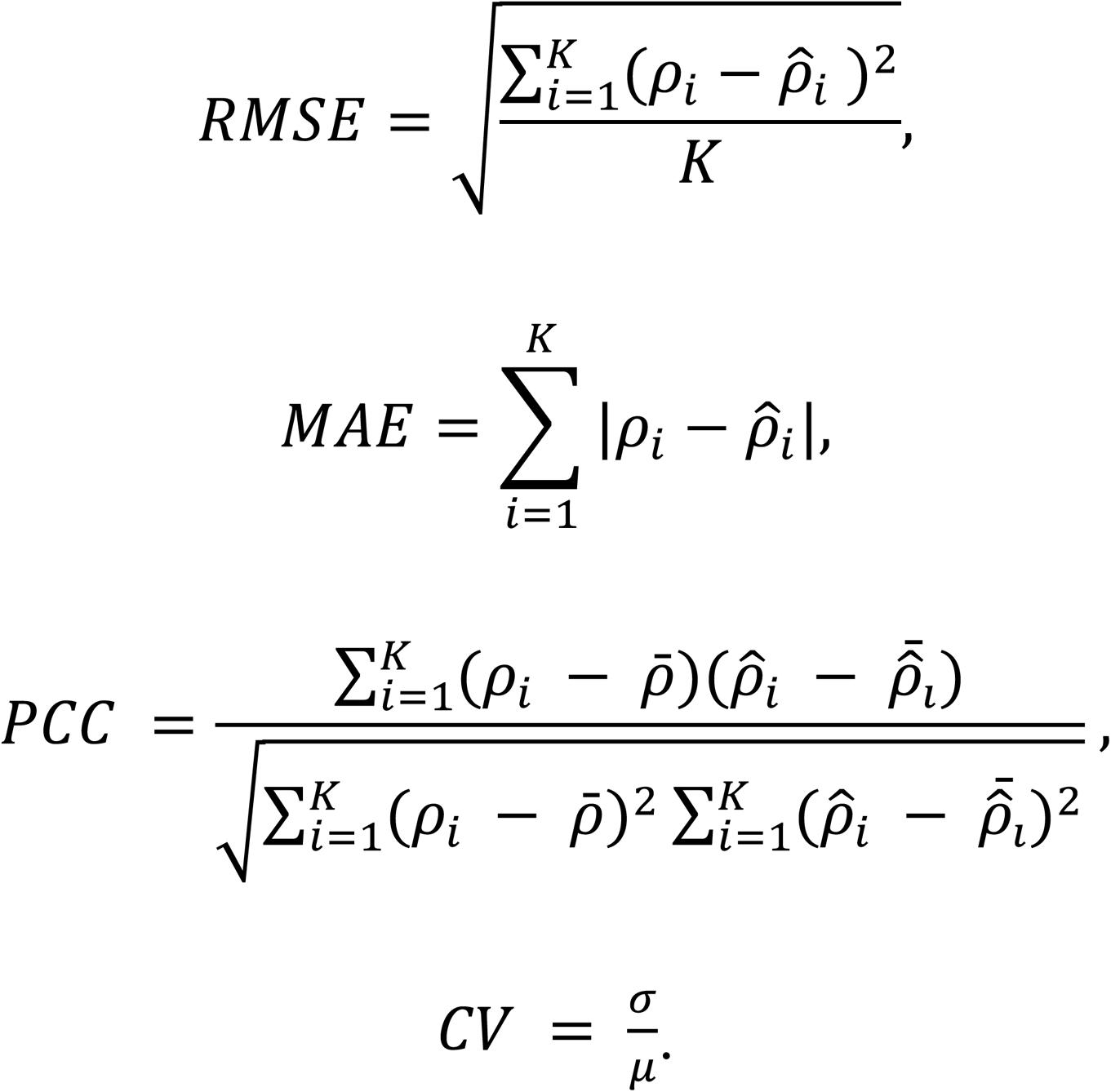

Here the predicted cell-type proportion is denoted as 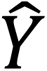 and ground truth cell-type proportion is denoted as *Y*.

### Spatial transcriptomics analysis

To evaluate the marker genes for cell types with spatial patterns, we consider the similarity between the gene-gene correlation from the reference scRNA-seq data and the gene-gene correlation from spatial transcriptomic data, inspired by this work ^58^. Since these two datasets come from the same tissue, the gene-gene correlation should also be preserved. We denote the gene expression profiles after marker genes selection as 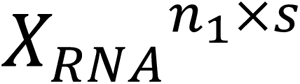 and 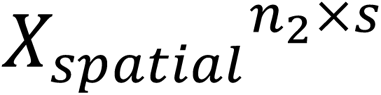. Using Pearson correlation, we can compute the gene-gene correlation matrices as 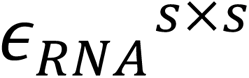 and 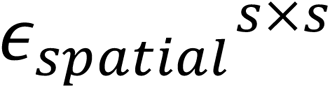. We calculate the Pearson correlation between these two matrices after flattening. A higher correlation represents a better marker gene set.

### Explanation of baseline models

In the benchmarking process for cell-type annotation and feature extraction, we consider five methods (settings): COSG, STG, scGeneFit, NS-Forest, and expert markers. The order of these methods (settings) is random.

COSG first defines an anchor gene based on its expression pattern for each cell type, then it utilizes a score defined based on cosine similarity between the anchor gene and the target gene to select marker genes.

STG is a method for general feature selection. STG learns a neural network with feature gates to automatically select important features based on optimizing the neural network. The number of important features is determined by the threshold to construct the gates.

scGeneFit treats the marker gene selection task as a projection problem. It selects marker genes by learning a projection to the lowest-dimensional space that maintains a clear separation between samples with different labels, thus generating a matrix with a smaller number of features. scGeneFit cannot select marker genes for different cell types.

NS-forest is a method for marker gene selection based on Random Forest (RF), feature ranking, Decision Tree (DT), and marker combinations. It first utilizes a RF model to learn the genes that are highly related to the cell types, and then filters the gene sets by ranking the binary score. After having the top binary gene list, NS-forest utilizes a DT model to assign these top binary genes to different cell types. By evaluating the combination of these marker genes based on the F-beta score, NS-forest can select the marker genes group with the highest F-beta score for each cell type.

Expert markers represent the marker genes of different cell types discovered by experts or biological experiments.

In the marker gene explanation section, we consider two methods: COSG and MetaMarker. The order of these methods is random.

MetaMarker is an explainable method for marker gene selection based on the trade-off between area-under-curve (AUC) score and fold change (FC) score. Metamarker prefers selecting a gene with a high expression rank and a high FC score in a given cell type as its marker gene.

In the analysis of marker gene selection for spatial transcriptomics, we consider five methods: COSG, STG, scGeneFit, NS-Forest, and scMAGS.

scMAGS is a method for transferring the marker gene information from scRNA-seq datasets to spatial transcriptomic datasets from the same tissue. It first transfers the scRNA-seq count matrix to an expression rate matrix and an average expression matrix, then for every cell type, scMAGS sets a threshold based on the expression rate matrix and only considers the genes above the threshold. scMAGS also defines a dispersion score by comparing the average expression value of the target gene in the given cell type with the average expression in other cell types, and finally selects genes with high dispersion rates.

In the analysis of deconvolution for bulk RNA-seq, we consider three methods: CIBERSORTx, MuSiC, and NNLS. The order of these methods is random.

CIBERSORTx is a method that employs machine learning to infer cell-type-specific gene expression profiles without physical cell isolation based on support vector regression.

MuSiC utilizes the weighted non-negative least squares algorithm to construct a hierarchical clustering tree, which gives each gene different scores. Then it calculates the proportion of clusters then determines the proportion of cell types.

NNLS uses the non-negative least squares regression to deconvolve a bulk sample.

### Experiment design

For scRNA-seq datasets, we followed the pre-processing guidelines from Scanpy ^24^. We filtered the MT-genes of the dataset based on raw counts followed by normalization and *log*(1 + *x*) transformation. After finishing the filtering and normalization steps, we ran marker gene selection methods after selecting the top 2000 HVGs.

For the bulk-seq datasets, we followed the pre-processing guidelines from CIBERSORT-X ^38^. We normalized the raw data. After selecting the maker genes based on scRNA-seq, we used our own pipeline to generate the deconvolution results.

For spatial transcriptomic datasets, we followed the pre-processing guidelines from Squidpy ^59^. After finishing the filtering and normalization steps, we run marker gene selection methods after selecting HVGs for the 10x Visium dataset and the human breast dataset (Xenium dataset). After selecting the maker genes, we use our own pipeline to generate the deconvolution results for the 10x Visium dataset.

## Data availability

We summarize the sources and statistics of all the datasets we used in Supplementary file 3. All the public datasets can be accessed based on the links in this file. The AD-HC dataset will be made publicly accessible upon the publication of this work.

## Reproductivity and Codes availability

We relied on Yale High-performance Computing Center (YCRC) and utilized one NVIDIA GTX1080Ti GPU with up to 30 GB RAM for model training. The codes of CosGeneGate can be found at https://github.com/VivLon/CosGeneGate. We follow the MIT license for usage.

## Acknowledgements

We thank Yingxin Lin and Yutaro Yamada for the suggestions and comments.

## Author contributions

T.L., Y.W., W.L. and H.Z. designed this study. W.L., Z.C. and T.L. ran all the experiments. T.L., W.L., Z.C., Y.W. and H.Z. wrote the manuscript. C.H., L.Z. and S.S. provided AD/HC datasets. H.Z. supervised this work.

## Competing interests

We do not have competing interests in this study.

## Ethics and Inclusion

Although CosGeneGate is not biased on gender, races, and other factors, the users are solely responsible for the content they generate with models in CosGeneGate, and there are no mechanisms in place for addressing harmful, unfaithful, biased, and toxic content disclosure. Any modifications of the models should be released under different version numbers to keep track of the original models related to this manuscript. The users must comply with the laws of the country in which they are located. The target of current CosGeneGate only serves for academic research. The users cannot use it for other purposes. Finally, we are not responsible for any effects produced by other users.

---

**Extended Data Fig. 1.**
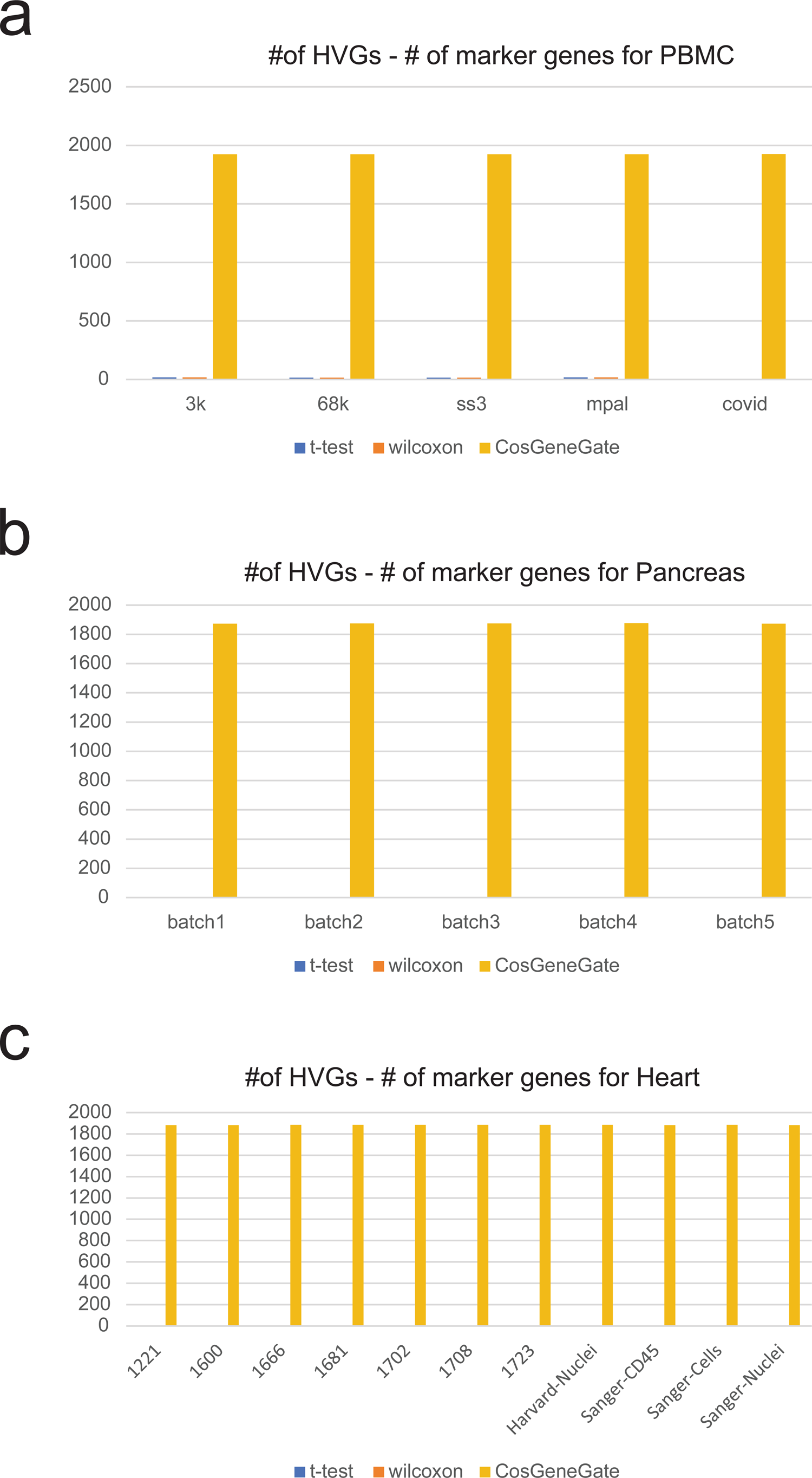
Difference between the number of HVGs and selected marker genes for different methods.

**Extended Data Fig. 2.**
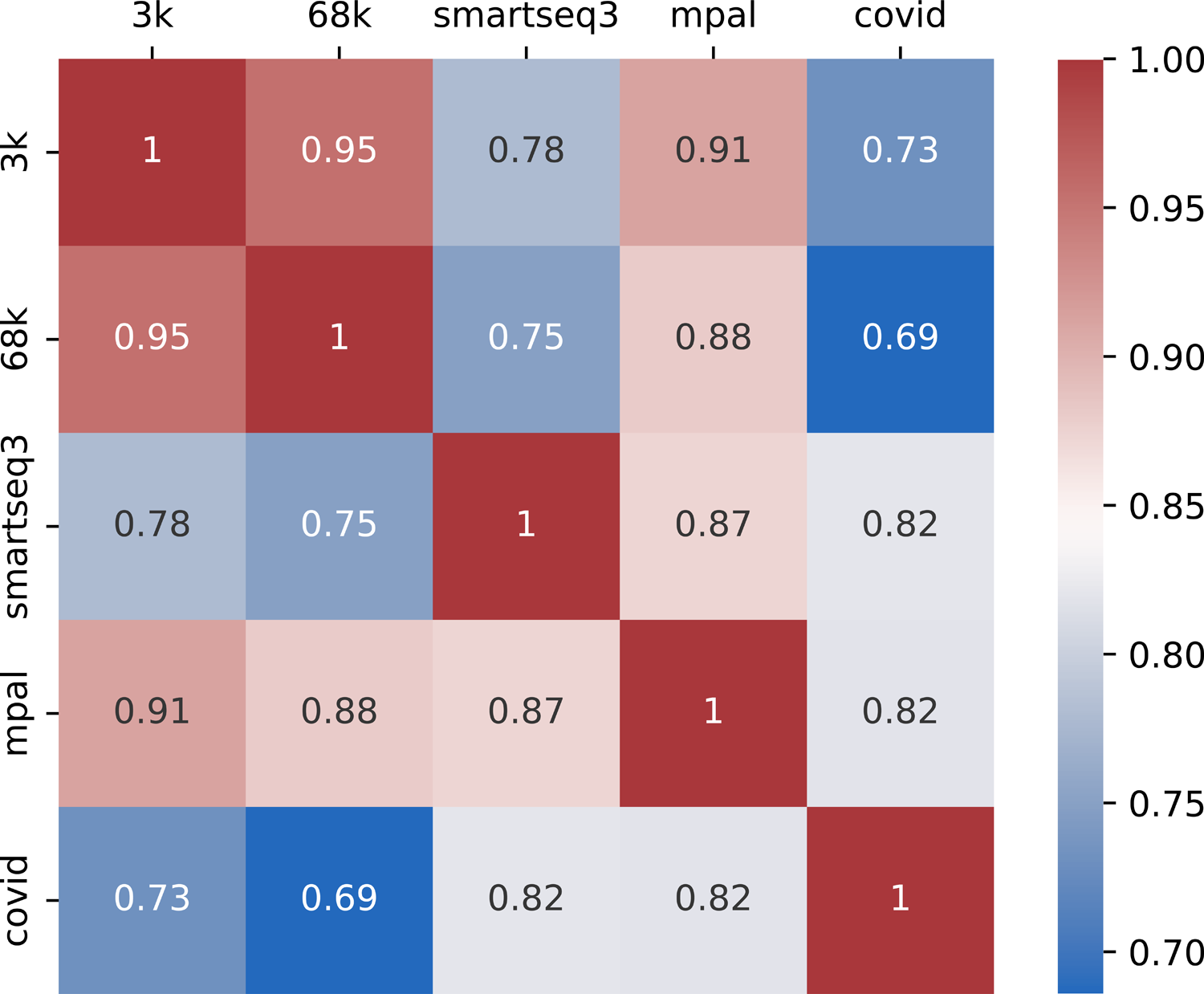
Cross-dataset cell-type similarity.

**Extended Data Fig. 3.**
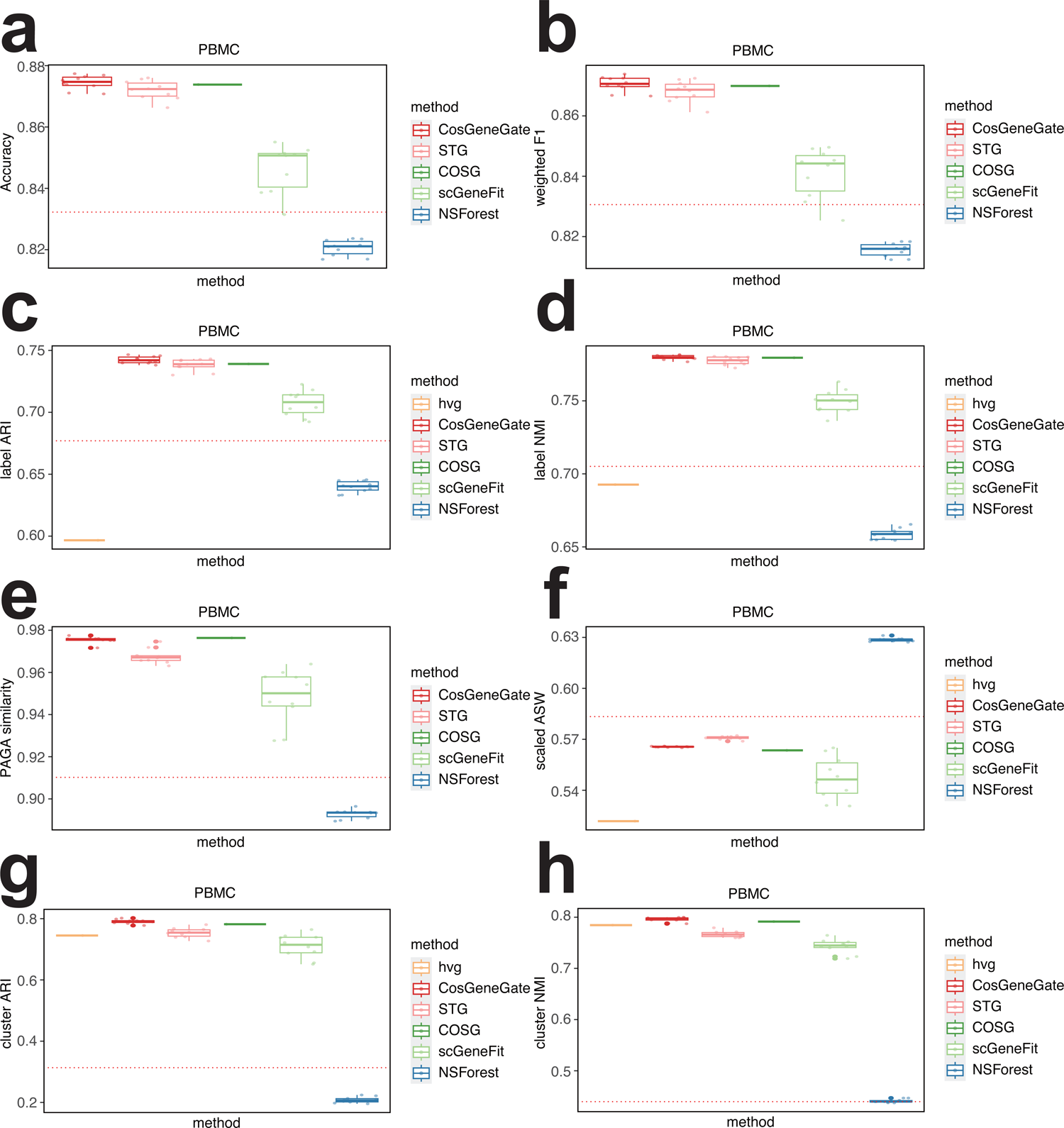
Boxplots for the annotation score and feature extraction score across different methods from PBMC. (a)-(g) are corresponding to metrics including Accuracy, weighted F1, label ARI, label NMI, PAGA similarity, scaled ASW, cluster ARI, cluster NMI.

**Extended Data Fig. 4.**
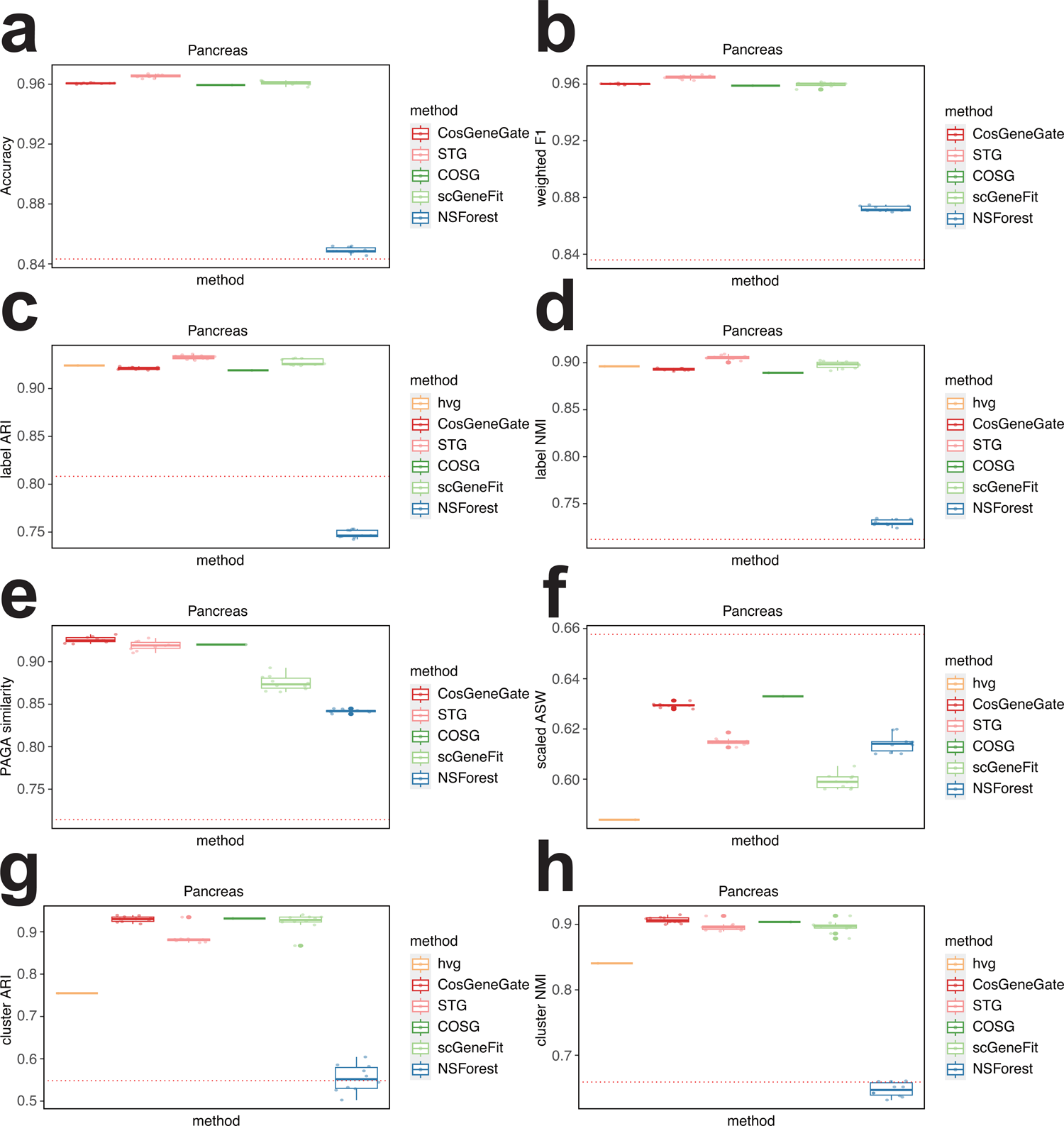
Boxplots for the annotation score and feature extraction score across different methods from Pancreas. (a)-(g) are corresponding to metrics including Accuracy, weighted F1, label ARI, label NMI, PAGA similarity, scaled ASW, cluster ARI, cluster NMI.

**Extended Data Fig. 5.**
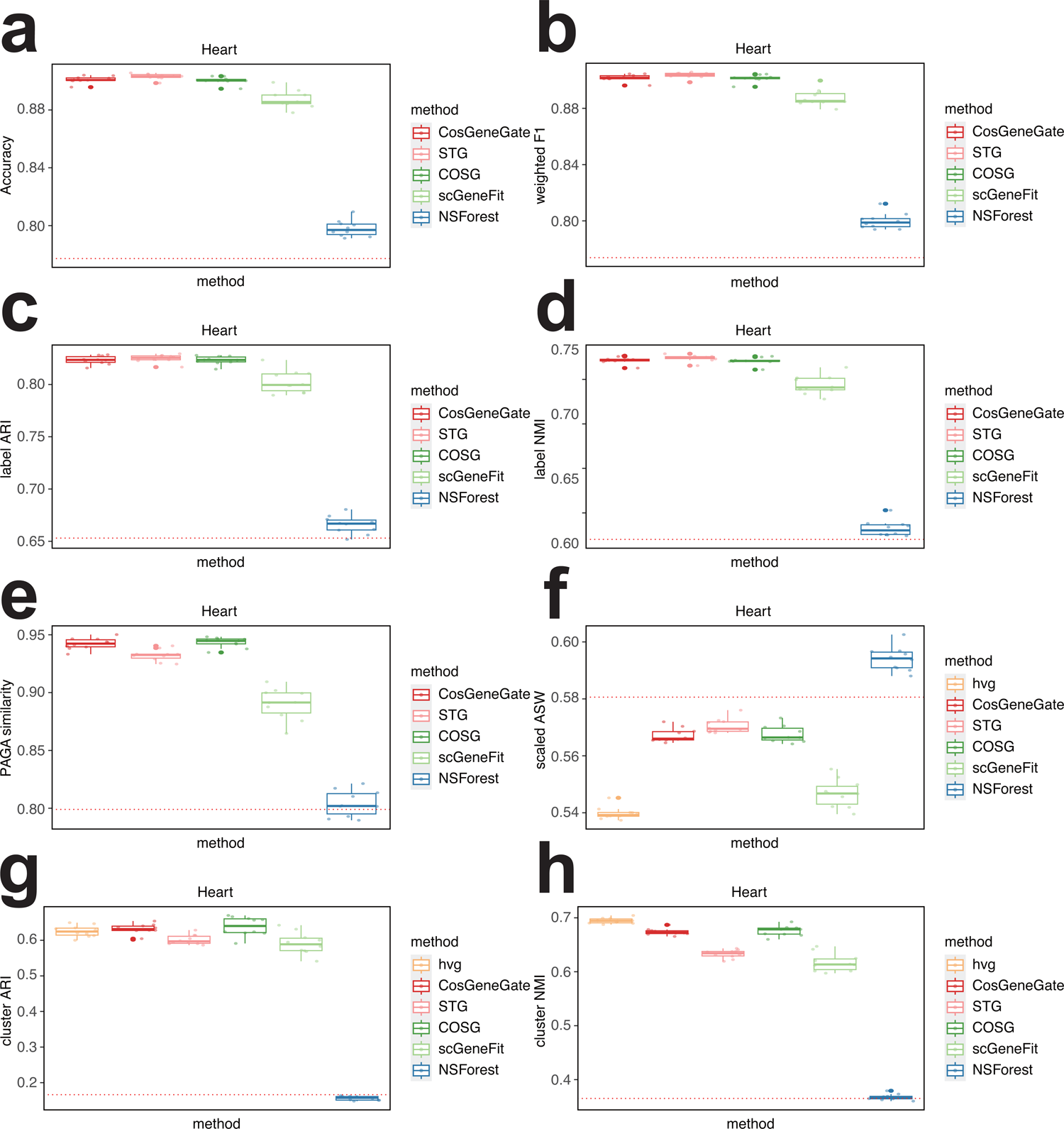
Boxplots for the annotation score and feature extraction score across different methods from Heart. (a)-(g) are corresponding to metrics including Accuracy, weighted F1, label ARI, label NMI, PAGA similarity, scaled ASW, cluster ARI, cluster NMI.

**Extended Data Fig. 6.**
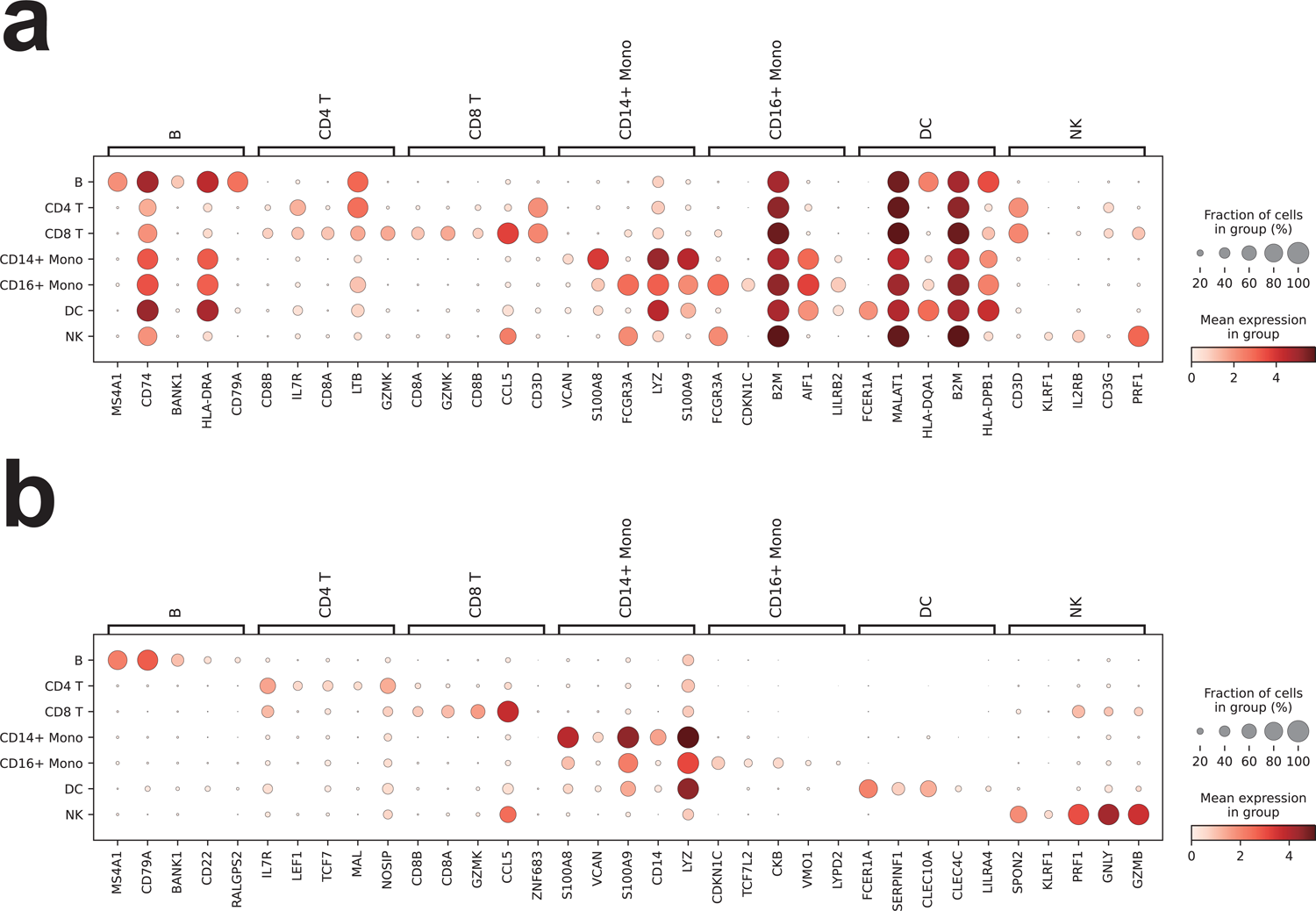
Expression profiles of STG and COSG markers. (a) Dotplot for markers from STG based on PBMC dataset. (b) Dotpot for markers from COSG based on PBMC dataset.

**Extended Data Fig. 7.**
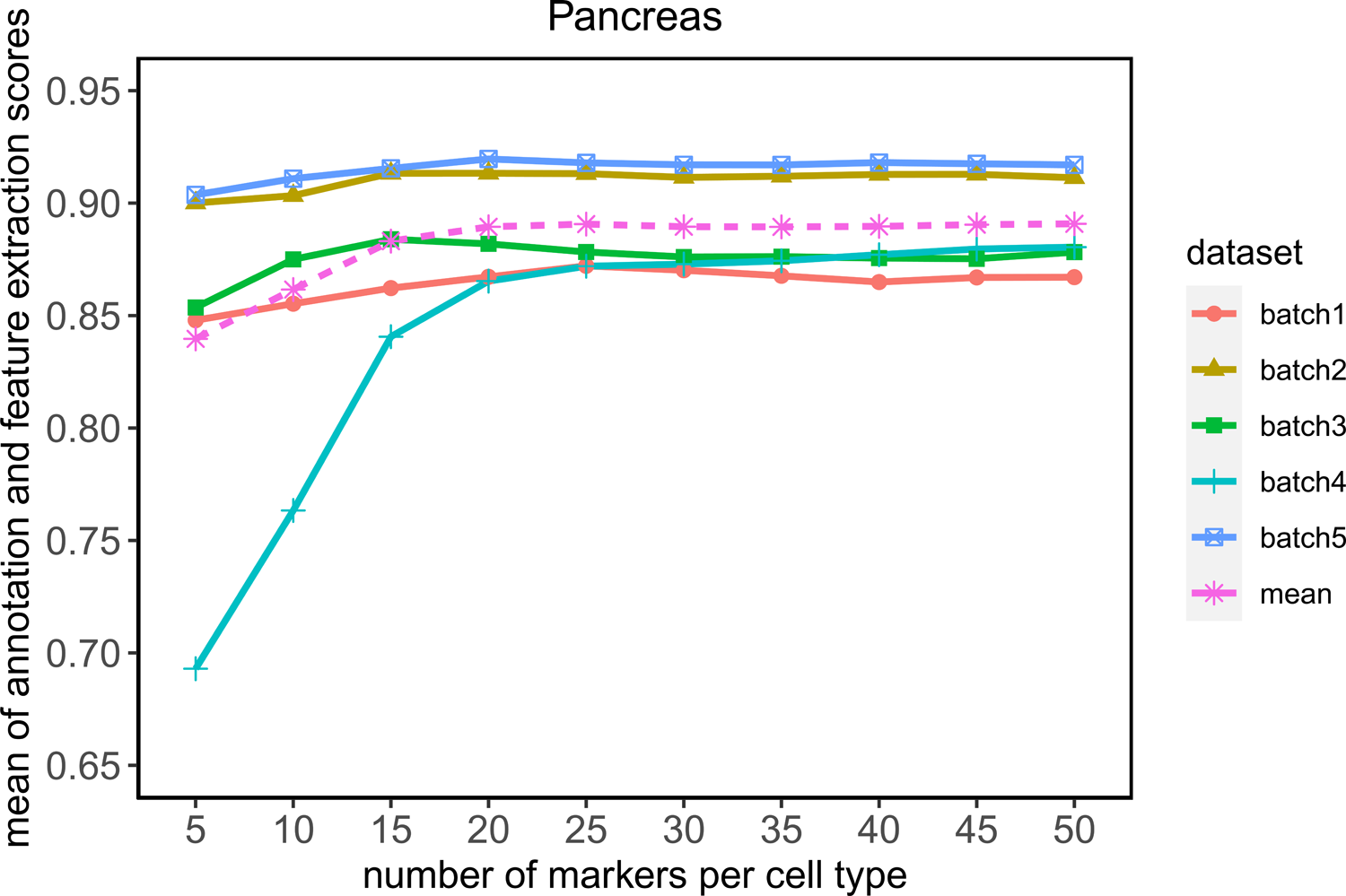
CosGeneGate hyper-parameter tuning for pancreas dataset.

**Extended Data Fig. 8.**
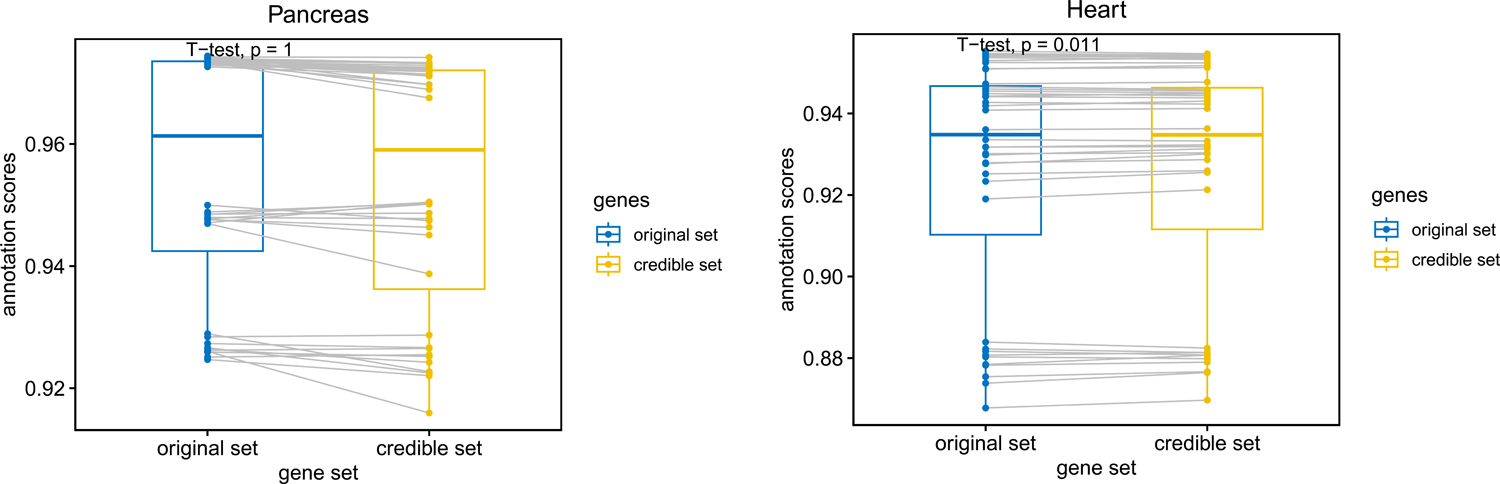
Comparisons between the original gene sets and gene sets after removing redundancy for the cell-type annotation accuracy.

**Extended Data Fig. 9.**
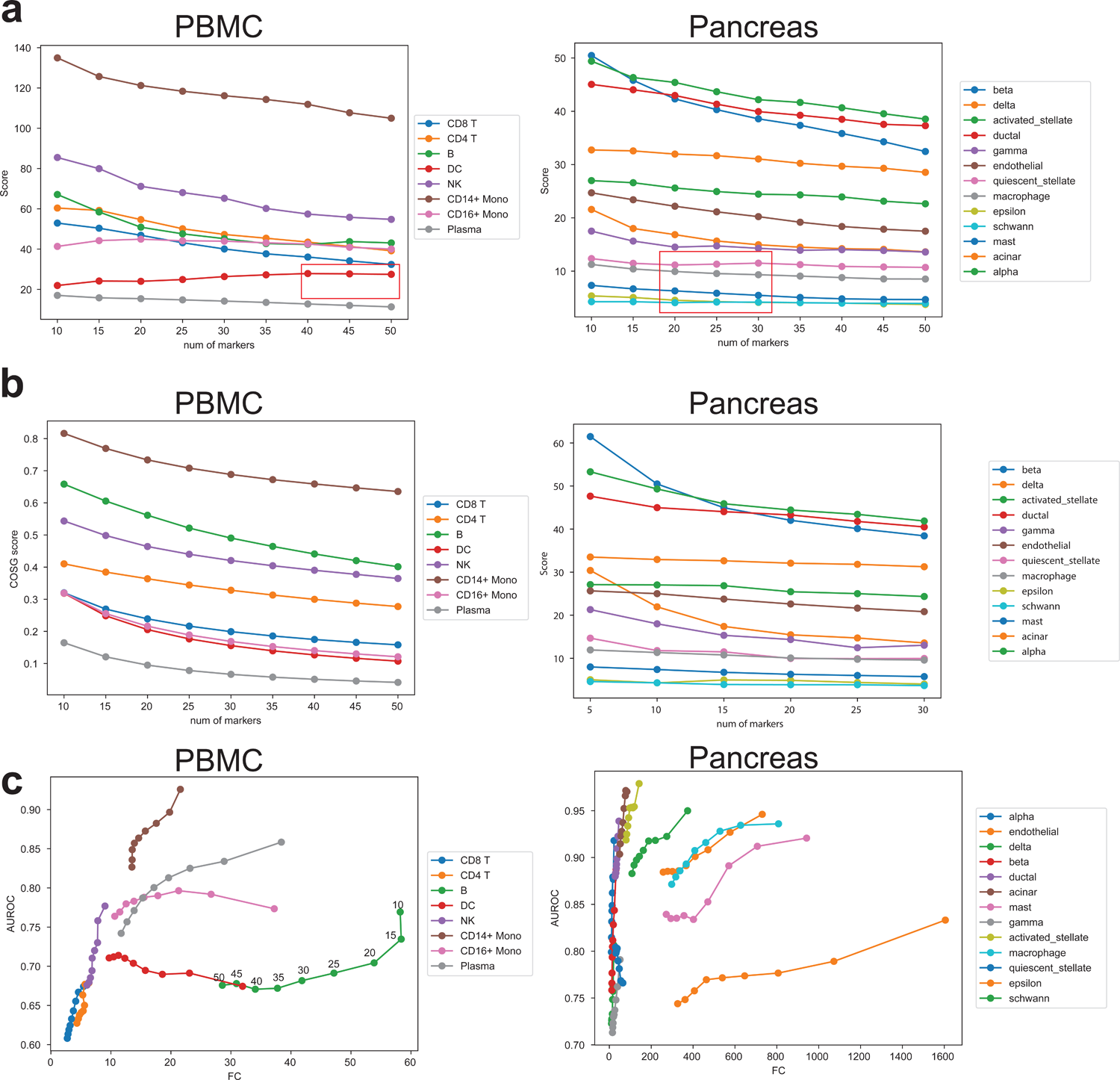
Explainability of selected markers based on different methods. (a) Relationship between the number of selected markers and the Wilcoxon score of CosGeneGate across different datasets. Different lines are colored by cell types. (b) Relationship between the number of selected markers and the Wilcoxon score of COSG across different datasets. Different lines are colored by cell types. (c) Relationship between Fold-change score and AUROC of MetaMarker across different datasets. Different lines are colored by cell types.

**Extended Data Fig. 10.**
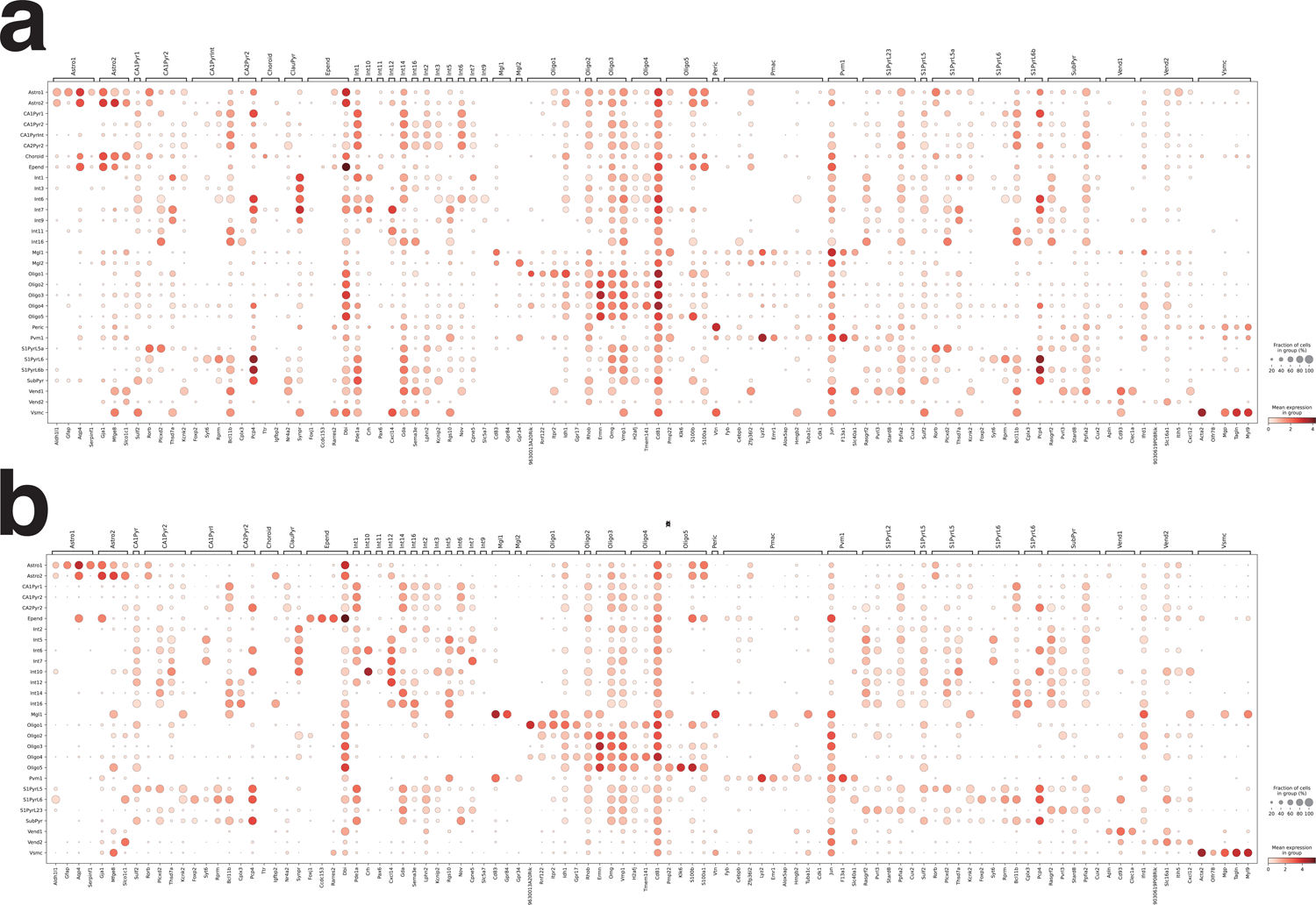
Full marker genes plots for cells with original cell types and refined cell types.

**Extended Data Fig. 11.**
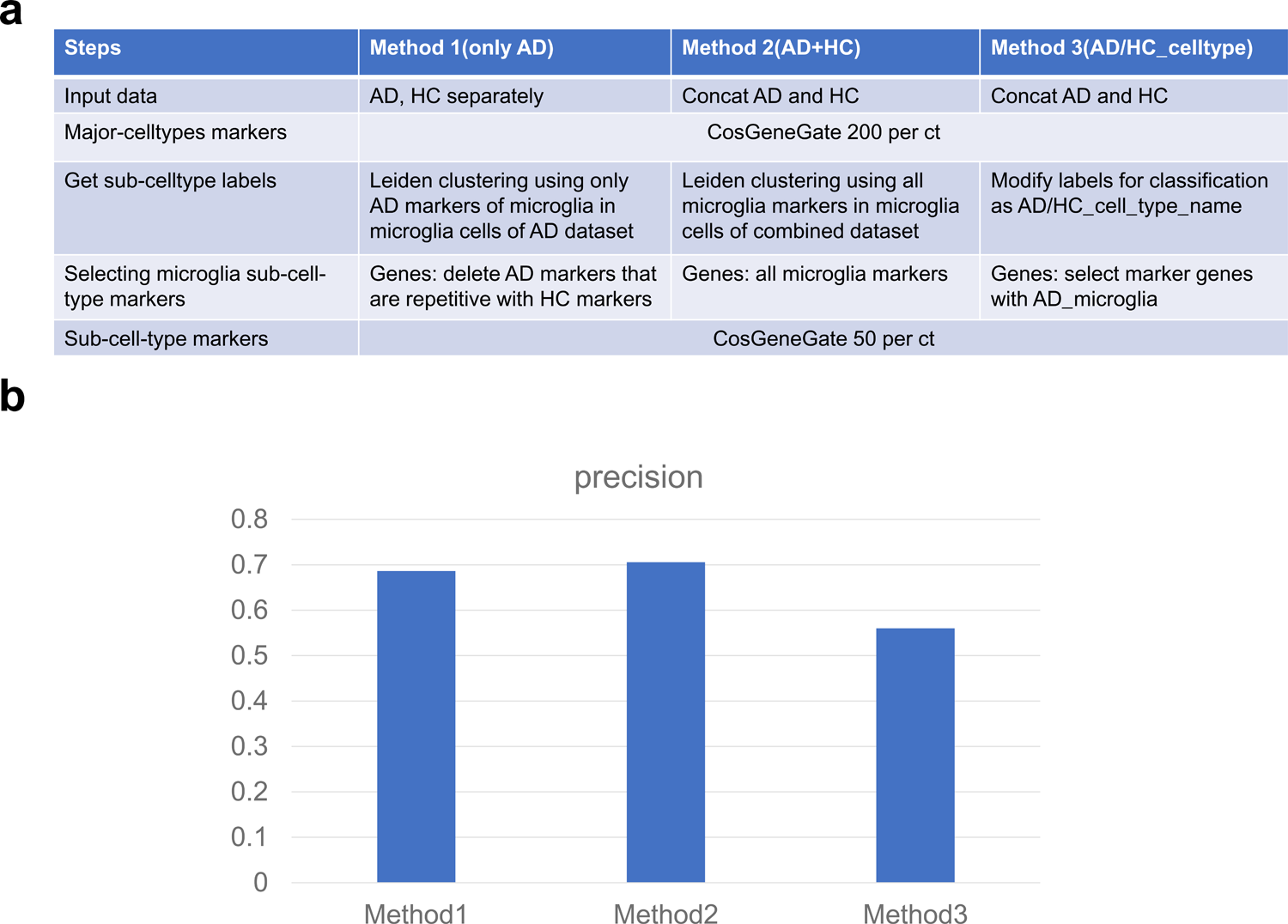
Comparison of three approaches to identify disease-specific marker genes for microglia cells. (a) A table to describe three different methods we used for AD-specific marker gene selection. (b) Precision scores across different methods.

**Extended Data Fig. 12.**
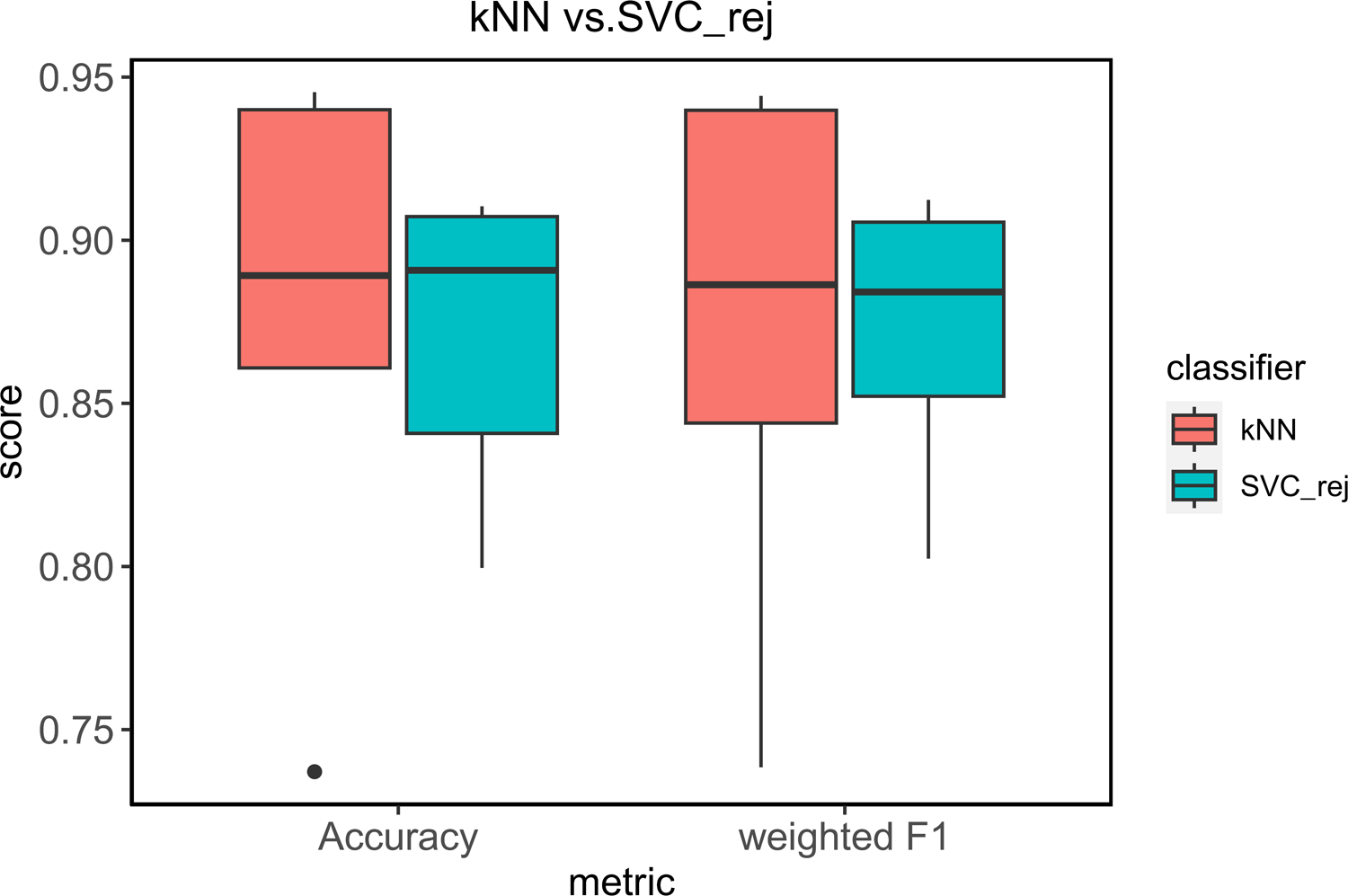
Annotation performance using different classifiers in the testing stage.

**Supplementary file 1: credible sets of major cell types in PBMC, Pancreas and Heart. Optimal gene sets are in red and the rest are in black.**

**Supplementary file 2: disease-specific marker genes information with GWAS support or biological research support.**

**Supplementary file 3: Dataset information.**

## Notes

### Competing Interest Statement

The authors have declared no competing interest.

## References

1. Han, X. et al. Construction of a human cell landscape at single-cell level. Nature 581, 303–309 (2020).

2. Saliba, A.-E., Westermann, A. J., Gorski, S. A. & Vogel, J. Single-cell RNA-seq: advances and future challenges. Nucleic Acids Res. 42, 8845–8860 (2014).

3. Mathys, H. et al. Single-cell transcriptomic analysis of Alzheimer’s disease. Nature 570, 332–337 (2019).

4. Stubbington, M. J. T., Rozenblatt-Rosen, O., Regev, A. & Teichmann, S. A. Single-cell transcriptomics to explore the immune system in health and disease. Science 358, 58–63 (2017).

5. Zhang, Z., et al. Signal Recovery in Single Cell Batch Integration. http://biorxiv.org/lookup/doi/10.1101/2023.05.05.539614 (2023) doi:10.1101/2023.05.05.539614.

6. Evrony, G. D., Hinch, A. G. & Luo, C. Applications of Single-Cell DNA Sequencing. Annu. Rev. Genomics Hum. Genet. 22, 171–197 (2021).

7. Hwang, B., Lee, J. H. & Bang, D. Single-cell RNA sequencing technologies and bioinformatics pipelines. Exp. Mol. Med. 50, 1–14 (2018).

8. Zheng, G. X. Y. et al. Massively parallel digital transcriptional profiling of single cells. Nat. Commun. 8, 14049 (2017).

9. Stoeckius, M. et al. Simultaneous epitope and transcriptome measurement in single cells. Nat. Methods 14, 865–868 (2017).

10. Cusanovich, D. A. et al. Multiplex single-cell profiling of chromatin accessibility by combinatorial cellular indexing. Science 348, 910–914 (2015).

11. Chen, M. et al. Differentiation of isomeric methylanilines by imidization and gas chromatography/mass spectrometry analysis. Rapid Commun. Mass Spectrom. 32, 342–348 (2018).

12. Luo, C. et al. Single-cell methylomes identify neuronal subtypes and regulatory elements in mammalian cortex. Science 357, 600–604 (2017).

13. Mulqueen, R. M. et al. Highly scalable generation of DNA methylation profiles in single cells. Nat. Biotechnol. 36, 428–431 (2018).

14. Zhang, M. et al. Spatially resolved cell atlas of the mouse primary motor cortex by MERFISH. Nature 598, 137–143 (2021).

15. Flynn, E., Almonte-Loya, A. & Fragiadakis, G. K. Single-Cell Multiomics. Annu. Rev. Biomed. Data Sci. 6, 313–337 (2023).

16. Fleck, J. S., Camp, J. G. & Treutlein, B. What is a cell type? Science 381, 733–734 (2023).

17. Yu, L., Wu, Y., Dunn, J. F. & Murari, K. In-vivo monitoring of tissue oxygen saturation in deep brain structures using a single fiber optical system. Biomed. Opt. Express 7, 4685 (2016).

18. Litviňuková, M. et al. Cells of the adult human heart. Nature 588, 466–472 (2020).

19. Yang, P., Huang, H. & Liu, C. Feature selection revisited in the single-cell era. Genome Biol. 22, 321 (2021).

20. Giladi, A. et al. Single-cell characterization of haematopoietic progenitors and their trajectories in homeostasis and perturbed haematopoiesis. Nat. Cell Biol. 20, 836–846 (2018).

21. Gómez-Chávez, F. et al. Maternal Immune Response During Pregnancy and Vertical Transmission in Human Toxoplasmosis. Front. Immunol. 10, 285 (2019).

22. Domínguez Conde, C., et al. Cross-tissue immune cell analysis reveals tissue-specific features in humans. Science 376, eabl5197 (2022).

23. Stuart, T. et al. Comprehensive Integration of Single-Cell Data. Cell 177, 1888–1902.e21 (2019).

24. Wolf, F. A., Angerer, P. & Theis, F. J. SCANPY: large-scale single-cell gene expression data analysis. Genome Biol. 19, 15 (2018).

25. Blondel, V. D., Guillaume, J.-L., Lambiotte, R. & Lefebvre, E. Fast unfolding of communities in large networks. J. Stat. Mech. Theory Exp. 2008, P10008 (2008).

26. Traag, V. A., Waltman, L. & Van Eck, N. J. From Louvain to Leiden: guaranteeing well-connected communities. Sci. Rep. 9, 5233 (2019).

27. Aevermann, B. et al. A machine learning method for the discovery of minimum marker gene combinations for cell type identification from single-cell RNA sequencing. Genome Res. 31, 1767–1780 (2021).

28. Dumitrascu, B., Villar, S., Mixon, D. G. & Engelhardt, B. E. Optimal marker gene selection for cell type discrimination in single cell analyses. Nat. Commun. 12, 1186 (2021).

29. Pullin, J. M. & McCarthy, D. J. A comparison of marker gene selection methods for single-cell RNA sequencing data. Genome Biol. 25, 56 (2024).

30. Yamada, Y., Lindenbaum, O., Negahban, S. & Kluger, Y. Feature Selection using Stochastic Gates. in Proceedings of the 37th International Conference on Machine Learning (eds. III, H. D. & Singh, A.) vol. 119 10648–10659 (PMLR, 2020).

31. Dai, M., Pei, X. & Wang, X.-J. Accurate and fast cell marker gene identification with COSG. Brief. Bioinform. 23, bbab579 (2022).

32. Wang, Y., Liu, T. & Zhao, H. ResPAN: a powerful batch correction model for scRNA-seq data through residual adversarial networks. Bioinformatics 38, 3942–3949 (2022).

33. Granja, J. M. et al. Single-cell multiomic analysis identifies regulatory programs in mixed-phenotype acute leukemia. Nat. Biotechnol. 37, 1458–1465 (2019).

34. Wilk, A. J. et al. Multi-omic profiling reveals widespread dysregulation of innate immunity and hematopoiesis in COVID-19. J. Exp. Med. 218, e20210582 (2021).

35. Tran, H. T. N. et al. A benchmark of batch-effect correction methods for single-cell RNA sequencing data. Genome Biol. 21, 12 (2020).

36. Wolf, F. A. et al. PAGA: graph abstraction reconciles clustering with trajectory inference through a topology preserving map of single cells. Genome Biol. 20, 59 (2019).

37. Fischer, S. & Gillis, J. How many markers are needed to robustly determine a cell’s type? iScience 24, (2021).

38. Newman, A. M. et al. Determining cell type abundance and expression from bulk tissues with digital cytometry. Nat. Biotechnol. 37, 773–782 (2019).

39. Wang, X., Park, J., Susztak, K., Zhang, N. R. & Li, M. Bulk tissue cell type deconvolution with multi-subject single-cell expression reference. Nat. Commun. 10, 380 (2019).

40. Danaher, P. et al. Advances in mixed cell deconvolution enable quantification of cell types in spatial transcriptomic data. Nat. Commun. 13, 385 (2022).

41. Dong, M. et al. SCDC: bulk gene expression deconvolution by multiple single-cell RNA sequencing references. Brief. Bioinform. 22, 416–427 (2021).

42. Biancalani, T. et al. Deep learning and alignment of spatially resolved single-cell transcriptomes with Tangram. Nat. Methods 18, 1352–1362 (2021).

43. Lin, Y. et al. scClassify: sample size estimation and multiscale classification of cells using single and multiple reference. Mol. Syst. Biol. 16, e9389 (2020).

44. Baran, Y. & Doğan, B. scMAGS: Marker gene selection from scRNA-seq data for spatial transcriptomics studies. Comput. Biol. Med. 155, 106634 (2023).

45. Zeisel, A. et al. Cell types in the mouse cortex and hippocampus revealed by single-cell RNA-seq. Science 347, 1138–1142 (2015).

46. Hansen, D. V., Hanson, J. E. & Sheng, M. Microglia in Alzheimer’s disease. J. Cell Biol. 217, 459–472 (2018).

47. Keren-Shaul, H. et al. A Unique Microglia Type Associated with Restricting Development of Alzheimer’s Disease. Cell 169, 1276–1290.e17 (2017).

48. Wang, C. et al. The effects of microglia-associated neuroinflammation on Alzheimer’s disease. Front. Immunol. 14, 1117172 (2023).

49. Zhang, L., et al. Single-Cell Transcriptomic Atlas of Alzheimer’s Disease Middle Temporal Gyrus Reveals Region, Cell Type and Sex Specificity of Gene Expression with Novel Genetic Risk for MERTK in Female. http://medrxiv.org/lookup/doi/10.1101/2023.02.18.23286037 (2023) doi:10.1101/2023.02.18.23286037.

50. Yu, L. & Liu, H. Redundancy based feature selection for microarray data. In Proceedings of the tenth ACM SIGKDD international conference on Knowledge discovery and data mining 737–742 (ACM, Seattle WA USA, 2004). doi:10.1145/1014052.1014149.

51. Su, C. et al. Cell-type-specific co-expression inference from single cell RNA-sequencing data. Nat. Commun. 14, 4846 (2023).

52. MacArthur, J. et al. The new NHGRI-EBI Catalog of published genome-wide association studies (GWAS Catalog). Nucleic Acids Res. 45, D896–D901 (2017).

53. Abdelaal, T. et al. A comparison of automatic cell identification methods for single-cell RNA sequencing data. Genome Biol. 20, 194 (2019).

54. Pedregosa, F. et al. Scikit-learn: Machine Learning in Python. J. Mach. Learn. Res. 12, 2825–2830 (2011).

55. Luecken, M. D. et al. Benchmarking atlas-level data integration in single-cell genomics. Nat. Methods 19, 41–50 (2022).

56. Thomas, P. D. et al. PANTHER : Making genome-scale phylogenetics accessible to all. Protein Sci. 31, 8–22 (2022).

57. Roohani, Y., Huang, K. & Leskovec, J. Predicting transcriptional outcomes of novel multigene perturbations with GEARS. Nat. Biotechnol. (2023) doi:10.1038/s41587-023-01905-6.

58. Song, D. et al. scDesign3 generates realistic in silico data for multimodal single-cell and spatial omics. Nat. Biotechnol. 42, 247–252 (2024).

59. Palla, G. et al. Squidpy: a scalable framework for spatial omics analysis. Nat. Methods 19, 171–178 (2022).

